# N-terminal β-strand underpins biochemical specialization of an ATG8 isoform

**DOI:** 10.1101/453563

**Authors:** Erin K. Zess, Cassandra Jensen, Neftaly Cruz-Mireles, Juan Carlos De la Concepcion, Jan Sklenar, Richard Imre, Elisabeth Roitinger, Richard Hughes, Khaoula Belhaj, Karl Mechtler, Frank L.H. Menke, Tolga Bozkurt, Mark J. Banfield, Sophien Kamoun, Abbas Maqbool, Yasin F. Dagdas

## Abstract

ATG8 is a highly-conserved ubiquitin-like protein that modulates autophagy pathways by binding autophagic membranes and numerous proteins, including cargo receptors and core autophagy components. Throughout plant evolution, ATG8 has expanded from a single protein in algae to multiple isoforms in higher plants. However, the degree to which ATG8 isoforms have functionally specialized to bind distinct proteins remains unclear. Here, we describe a comprehensive protein-protein interaction resource, obtained using *in planta* immunoprecipitation followed by mass spectrometry, to define the potato ATG8 interactome. We discovered that ATG8 isoforms bind distinct sets of plant proteins with varying degrees of overlap. This prompted us to define the biochemical basis of ATG8 specialization by comparing two potato ATG8 isoforms using both *in vivo* protein interaction assays and *in vitro* quantitative binding affinity analyses. These experiments revealed that the N-terminal β-strand—and, in particular, a single amino acid polymorphism—underpins binding specificity to the substrate PexRD54 by shaping the hydrophobic pocket that accommodates this protein’s ATG8 interacting motif. Additional proteomics experiments indicated that the N-terminal β-strand shapes the ATG8 interactor profiles, defining interaction specificity with about 80 plant proteins. Our findings are consistent with the view that ATG8 isoforms comprise a layer of specificity in the regulation of selective autophagy pathways in plants.

## Introduction

Macroautophagy (hereafter autophagy) is a conserved cellular quality control pathway that removes unwanted self‐ and non-self macromolecules to maintain homeostasis in the face of physiological and environmental fluctuations [1,2]. There is a growing appreciation that autophagy is a highly selective process, with multiple layers of specificity defining the dynamics of uptake, sub-cellular trafficking, and turnover of autophagic substrates [3,4]. However, despite these advances, the molecular details of how various autophagy cargoes and components are recognized, recruited, and recycled remain to be fully elucidated [5,6]. In particular, our understanding of the molecular codes that define selective autophagy in plants is limited [7,8].

The autophagy machinery consists of around 40 autophagy related proteins (ATGs), which are highly conserved across eukaryotes [2,9]. Core ATG proteins mediate the repeated insertion of the ubiquitin-like protein ATG8 into the growing autophagosome membrane, culminating in the formation of a double-membraned vesicle. Generally, these mature autophagosomes are trafficked through the cell to fuse with the vacuole where autophagic cargoes are degraded and their building blocks—nucleic acids, amino acids, lipids and carbohydrates—are recycled and returned to the cytoplasm [10]. Autophagy can also participate in other processes besides the recycling of cellular components. For example, recent studies have implicated autophagy in unconventional secretion of cytosolic proteins [11,12], and susceptibility to invading pathogens [13,14].

ATG8 is a key player in the selective autophagy pathway. Besides being the major structural component of autophagosome membranes, ATG8 binds selective autophagy receptors and adaptors, as well as core ATG proteins, and autophagy-specific SNAREs (Soluble NSF Attachment protein Receptors) that mediate fusion of the autophagosome with the vacuole [10,15–17]. ATG8-interacting proteins often carry a conserved motif named the ATG8 Interacting Motif (AIM, also known as the LIR motif), which follows a W/Y/F-X-X-L/I/V consensus sequence and is typically surrounded by negatively charged residues [18]. ATG8 is composed of four α-helices and four β-strands, with a variable N-terminal extension that mediates the growth of the nascent autophagosome via fusion of ATG8-containing vesicles [19–21]. There are multiple ATG8 isoforms in metazoans that appear to be functionally specialized based on interactome analyses [22,23] and a few recent functional studies [24–28]. One emerging view is that ATG8 specialization could form a layer of specificity in selective autophagy pathways in addition to the autophagy cargo receptors. This could occur through ATG8 interaction with different sets of proteins [22,23,29], post-translational modifications, such as ubiquitination [26] and acetylation [30], and localization to different sub-cellular compartments [31]. However, despite these recent and consequential advances, ATG8 specificity has yet to be functionally studied in plants.

Phylogenetic analyses revealed that ATG8 has dramatically expanded throughout the evolution of land plants compared to algae, which only carries a single ATG8 [32]. ATG8 isoforms in land plants range from four in rice to twenty-two in oilseed rape [32]. Plant ATG8s group into two major clades: Clade I contains all of the algal and most angiosperm ATG8s, whereas Clade II is unique to land plants. Interestingly, ATG8s cluster in phylogenetic groups that are specific to particular plant families, indicating that ATG8s have diversified across plant taxa. For example, Solanaceae, the family that contains potato and Nicotiana *spp.*, have four well-supported monophyletic ATG8 clades that do not include ATG8s from other plant species, such as the model plant Arabidopsis [32]. A plausible hypothesis is that distinct ATG8 lineages have acquired specific polymorphisms throughout evolution that may have contributed to the diversification of selective autophagy pathways in plants [32]. Nonetheless, the degree to which these polymorphisms underlie ATG8 functional specialization remains unknown.

We recently showed that PexRD54, a secreted virulence protein from the Irish potato famine pathogen *Phytophthora infestans*, binds potato ATG8 isoform ATG8-2.2 (formerly ATG8CL, Clade I) via a canonical C-terminal AIM to modulate selective autophagy in host plant cells [33,34]. We solved the crystal structure of ATG8-2.2 in complex with a 5 amino acid peptide matching the C-terminal AIM of PexRD54, and showed that this AIM peptide docks within two hydrophobic pockets, W and L, similar to the yeast and metazoan ATG8-AIM complexes [34]. Interestingly, PexRD54 has a ten-fold higher binding affinity towards ATG8-2.2 compared to the potato Clade II ATG8-4 isoform (formerly ATG8IL) [33]. Additional *in planta* biochemical and cell biology assays further confirmed that PexRD54 has higher affinity to ATG8-2.2 compared to ATG8-4. For instance, PexRD54 increased accumulation of ATG8-2.2 autophagosomes and re-routed them towards the pathogen interface, but failed to significantly label and perturb ATG8-4 autophagosomes [33,35]. These findings indicate that these two potato ATG8 isoforms have different biochemical activities, and prompted us to further explore the hypothesis that plant ATG8 proteins are functionally specialized.

In this study, we used quantitative biochemistry and proteomics to explore the molecular basis of ATG8 specialization in solanaceous plants. First, we used immunoprecipitation followed by mass spectrometry analyses to determine that the six ATG8 potato isoforms bind distinct sets of plant proteins with varying degrees of overlap. Domain swaps and structure-guided mutagenesis experiments revealed that a recently derived polymorphism within the N-terminal β-strand of ATG8-4 underpins weak binding affinity towards PexRD54 and contributes to enhanced selectivity towards AIM-containing plant proteins. Our findings are a first step towards decoding the molecular signatures that define selective autophagy in plants. We propose that ATG8 specialization is an important specificity layer that determines the recruitment of cellular proteins to modulate selective autophagy pathways. In addition, the potato ATG8 interactome we produced reveals a number of novel proteins that are likely to play important roles in selective autophagy responses, thus serving as a valuable community resource and a foundation for future studies.

## Results

### Solanaceous ATG8 isoforms associate with distinct sets of proteins

Previous phylogenetic analyses revealed that the ATG8 family is expanded in plants and that ATG8 isoforms form well-supported family-specific clades, such as in the Solanaceae where ATG8s cluster into four major clades [32]. Potato encodes six ATG8 isoforms that cluster in these clades among close homologs from other solanaceous species (**Fig. 1a**; **Fig. S1**; **Fig. S2**). Based on sequence divergence (**Fig. S3**), we hypothesized that clade-specific ATG8s would interact with specific plant proteins and thus exhibit a degree of functional specialization.

To investigate this hypothesis, we determined the interactor profiles of all six potato ATG8s using immunoprecipitation (IP) coupled with mass spectrometry (MS) following transient expression in *Nicotiana benthamiana*. Our interactome analyses revealed 621 proteins that associated with at least one ATG8 isoform, but not with the empty vector control (**Table S1**). Consistent with our hypothesis, the potato ATG8s associated with unique complements of proteins with varying degrees of similarity, with ATG8-4 exhibiting the most selectivity (**Fig. 1b**; **Fig. S5**). The ATG8s showed differential interactions with a number of protein sets defined by cellular compartments (**Fig. 1c, Fig. S6**) and biological processes (**Fig. S7**), with the remaining sets common to all ATG8s. Within the interactome, we detected several core autophagy proteins and known ATG8-interacting proteins, including endogenous ATG8s, validating our IP-MS approach (**Table S1**; **Fig. S8**). We also captured a number of vesicle trafficking-related proteins, including Rab GTPases, Rab GTPase activating proteins, myosins, clathrin and coatomer subunits (**Table S1**) that were also identified in interactome studies using human ATG8 isoforms [22,23]. Excitingly, we found several hitherto unknown ATG8 associated proteins, including proteins that are predicted to localize to various organelles, making this dataset a useful community resource for future functional studies interrogating organelle recycling. Around 40% of all interactors are predicted to contain ATG8-interacting motifs (AIMs), of which a majority (73%) are conserved in either *Arabidopsis thaliana* or *Marchantia polymorpha*, suggesting the reliability of this dataset to further selective autophagy studies in plants (**Table S1**; **Table S2**).

Overall, the IP-MS screen indicated that solanaceous ATG8s have distinct interactor profiles, with ATG8-4 showing the most striking pattern. This prompted us to hypothesize that certain ATG8 domains or residues determine substrate binding specificity and underpin ATG8 specialization.

**Fig. 1.**
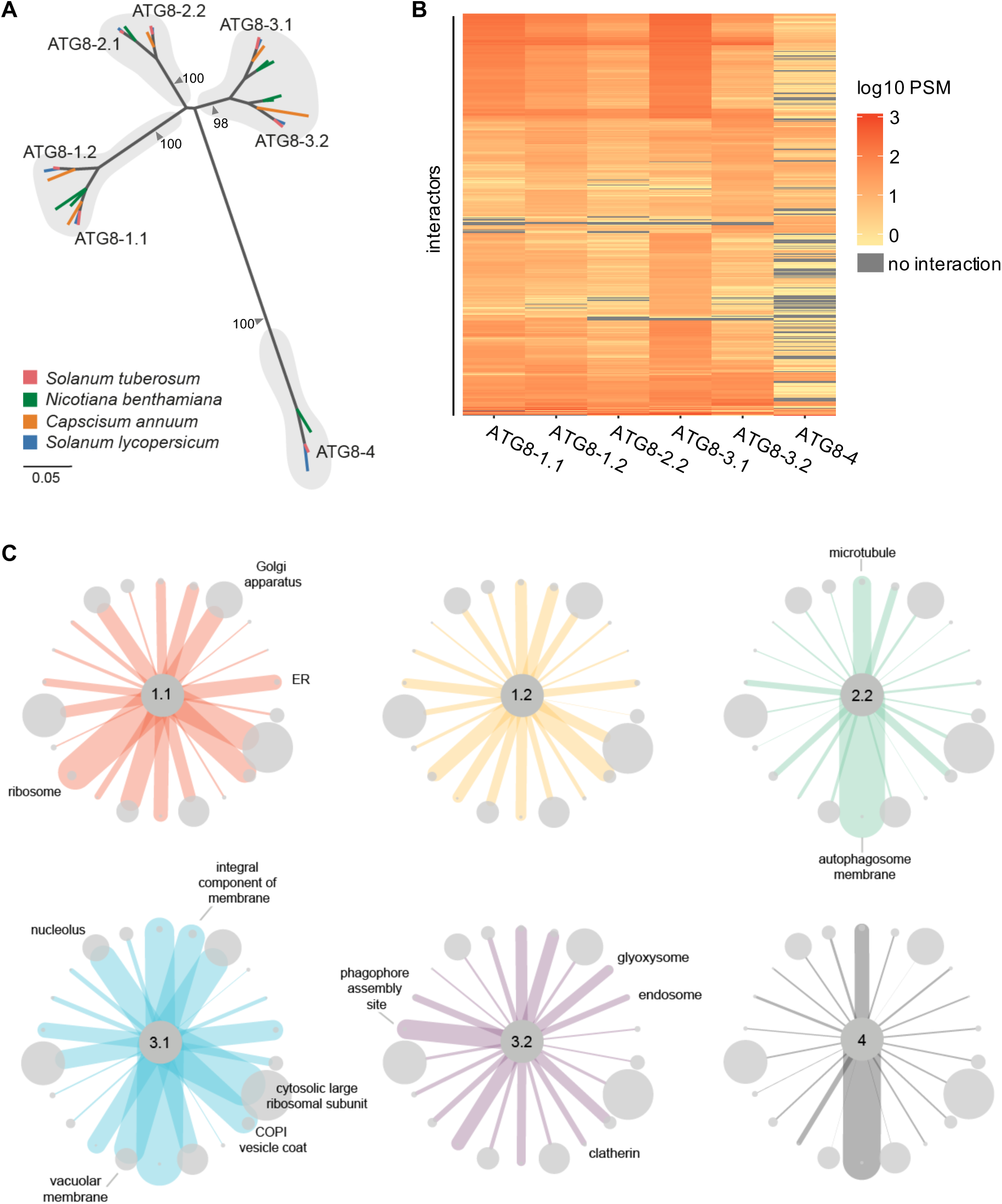
Solanaceous ATG8 isoforms have distinct plant protein interaction profiles. (a) Orthologous relationships between solanaceous ATG8 isoforms. Unrooted maximum-likelihood phylogenetic tree of 29 ATG8 isoforms with gray shading highlighting clades, and colors indicating plant species. The tree was calculated in MEGA7 [36] from a 369 nucleotide alignment (MUSCLE [37], codon-based). The bootstrap supports of the major nodes are indicated. The scale bar indicates the evolutionary distance based on nucleotide substitution rate, (b) Heatmap showing the interaction profiles of all ATG8 isoforms. The average PSM data was log10 normalized, and then used to construct a hierarchically clustered heatmap with the scale as shown, (c) Network representation of the interactions between ATG8s and protein groups defined by gene ontology (GO) cellular compartment annotations. Proteins were grouped based on the cellular compartment annotations, and a subset of groups were chosen for representation. The sizes of the nodes are scaled to the number of interactors in each respective group, and the edges are weighted to the average PSM values for all the interactors in each respective group for each ATG8. Nodes are labelled where the average PSM value is most differential compared to the other ATG8s. Figure S6 provides a graphical figure legend.

### ATG8 isoforms show differential binding to PexRD54

Our previous studies have shown that ATG8-2.2 and ATG8-4 have clear differences in binding affinity towards PexRD54. Therefore, we decided to use PexRD54 as a tool to dissect the structural elements within potato ATG8s that determine binding specificity to AIM-containing substrates. The potato ATG8s showed a range of association strengths with PexRD54 in *in planta* co-immunoprecipitation (Co-IP) experiments, indicating the capacity for ATG8s to selectively bind the same substrate (**Fig. 2a**). To quantify these affinity differences *in vitro*, we expressed and purified all potato ATG8 isoforms from *E. coli* (**Fig. S9a-b**). We then tested these ATG8 isoforms for binding with the PexRD54 AIM peptide using isothermal titration calorimetry (ITC) experiments. The ATG8 isoforms interacted with the AIM peptide with varying degrees of strength ranging from 29 nM to 287 nM (**Fig. 2b; Fig. S9c**). We then performed an additional biological replicate of our ITC experiments, where we obtained similar binding affinities (**Fig. 2c.**; **Fig. S9c-d**). Overall, ITC affinity measurements correlated with the trends in binding strength observed in the Co-IP experiments, where we used the full length PexRD54 protein, indicating that the AIM peptide alone is sufficient to recapitulate ATG8 binding specificities. In both methods, ATG8-2.2 and ATG8-4 displayed the greatest difference in PexRD54 binding strength, in a range that is consistent with our previous findings [33,34] (**Fig. 2c**).

### The first β-strand of ATG8 underpins discriminatory binding to PexRD54

To further investigate the ATG8 structural features that underpin discriminatory binding to PexRD54, and by proxy AIM-containing substrates, we generated chimeric proteins between ATG8-2.2 and ATG8-4 (**Fig. 3a**). We sequentially replaced ATG8-4 domains, the weakest PexRD54 interactor, with the corresponding domains from ATG8-2.2, the strongest interactor. Altogether we obtained a suite of eight ATG8 chimeras (Swaps 1- 8) and assayed these for gain-of-binding to PexRD54 (**Fig. 3a**). In Co-IP experiments, ATG8 chimera swap 3 (ATG8-4-S3), encompassing the first β-strand (β1), restored binding to PexRD54 (**Fig. 3b**). ATG8 chimera swap 1 (ATG8-4-S1) showed weak interaction with PexRD54, although this result was not consistent across replicates (**Fig. 3b**).

To validate these results using quantitative assays, we purified ATG8-4-S1 and ATG8-4-S3 from *E. coli* and tested for binding to both PexRD54 full length proteins and the AIM-peptide (**Fig. S10a-b**). Consistent with the Co-IP results, isothermal titration calorimetry measurements showed that the chimera ATG8-4-S3 bound to PexRD54 full length protein as well as the AIM-peptide with a similar affinity as ATG8-2.2, whereas ATG8-4-S1 bound weakly (**Fig. 3c**). We repeated these experiments and obtained similar results (**Fig. 3d; Fig. S10c-e**). Together, these results indicate that the ATG8 region encompassing the first β-strand (β1) is crucial for binding to PexRD54, specifically via this substrate’s AIM.

**Fig. 2.**
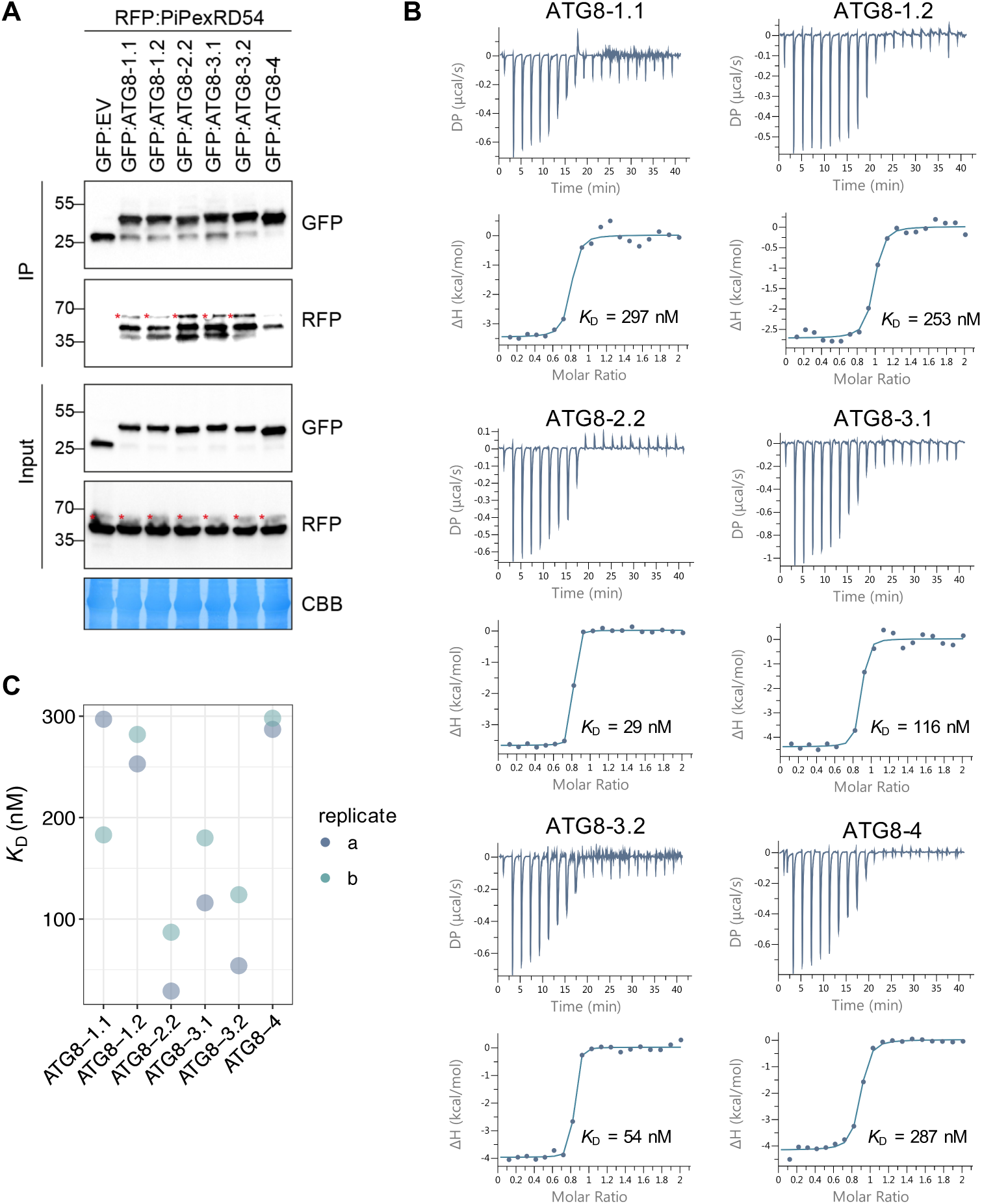
ATG8 isoforms show differential binding to PexRD54. (a) Co-Immunoprecipitation experiment between PexRD54 and all potato ATG8 isoforms. RFP:PiPexRD54 was transiently co-expressed with GFP:EV and all potato GFP:ATG8s. Immunoprecipitates (IPs) were obtained with anti-GFP antiserum and total protein extracts were immunoblotted with appropriate antisera (listed on the right). Stars indicate expected band sizes. (b) The binding affinities between ATG8 isoforms and the PexRD54 AIM peptide were determined using isothermal titration calorimetry (ITC). The top panels show heat differences upon injection of peptide ligands and lower panels show integrated heats of injection (•) and the best fit (solid line) to a single site binding model using MicroCal PEAQ-ITC analysis software (c) Chart summarizing the *K*_D_ value for each interaction across two replicates and highlighting variation within the replicates.

**Fig. 3.**
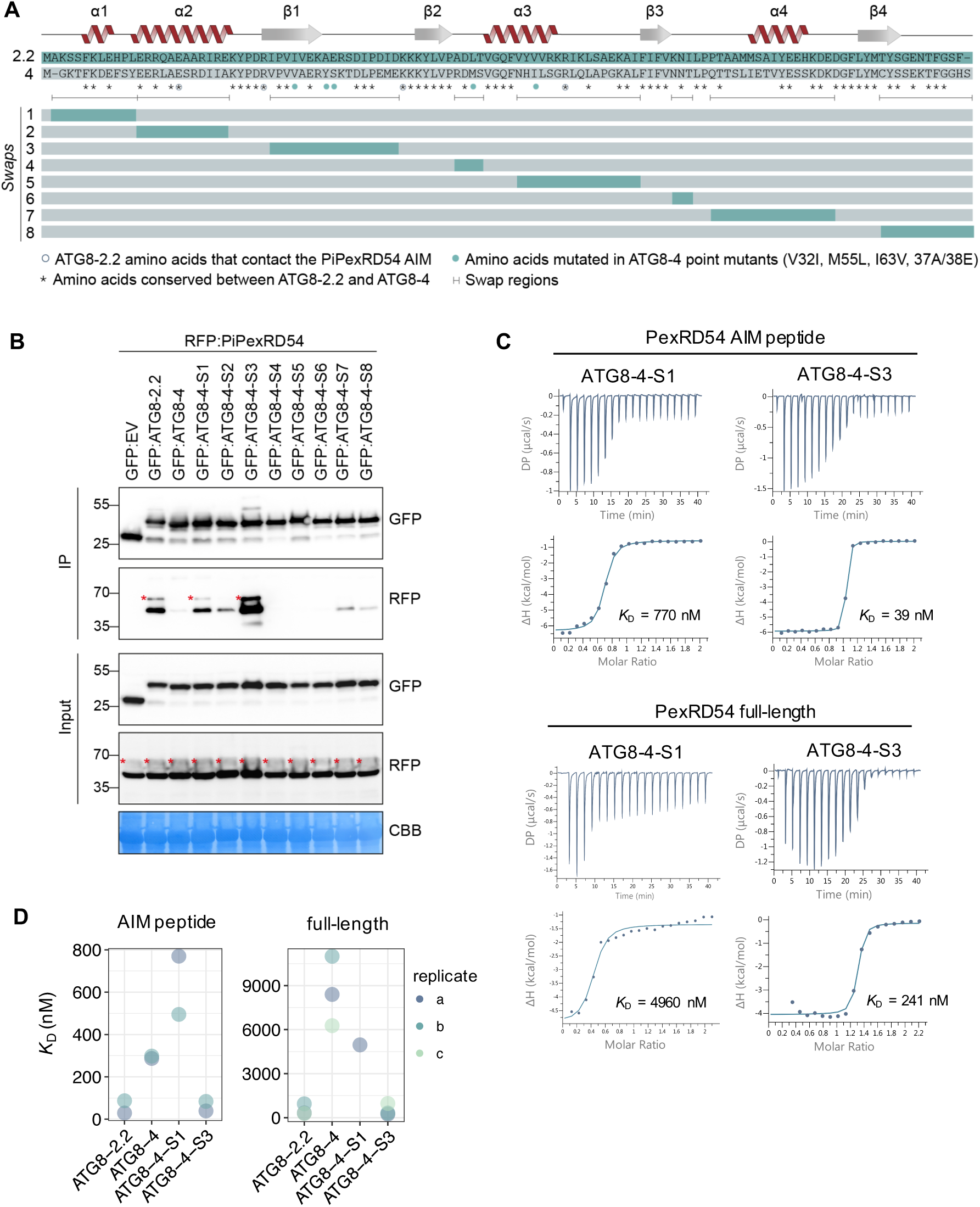
The ATG8 region surrounding the first β-strand is responsible for discriminatory binding to PexRD54. (a) Schematic showing the ATG8 swap chimeras and point mutants. The amino acid sequences of ATG8-2.2 and ATG8-4 are aligned, with the protein model above corresponding to the ATG8-2.2 structure. The brackets beneath the alignment indicate the boundaries of the swaps, with the color-coded rectangles below showing the chimeras made for each swap. The symbols beneath the alignment correspond to different features of the sequences: (i) the asterisks (*) mark conserved residues between ATG8-2.2 and ATG8-4, (ii) open circles mark residues that directly contact the PiPexRD54 AIM, and (iii) the filled circles mark the ATG8-4 residues used in structure guided mutagenesis experiments to match ATG8-2.2 (V32I, M55L, 163V, 37A/38E). (b) Co-immunoprecipitation experiment between PexRD54 and all ATG8 swap chimeras. RFP:PiPexRD54 was transiently co-expressed with the controls GFP:EV, GFP:ATG8-2.2, and GFP:ATG8-4, and all of the GFP:ATG8 swap chimeras (Swaps 1-8). Immunoprecipitates (IPs) were obtained with anti-GFP antibody and total protein extracts were immunoblotted with appropriate antibodies (listed on the right). Stars indicate expected band sizes, (c) The binding affinities of ATG8-4-S1 and ATG8-4-S3 towards PexRD54 AIM peptide and full-length PexRD54 were determined using isothermal titration calorimetry (ITC). The top panels show heat differences upon interaction and lower panels show integrated heats of injection (•) and the best fit (solid line) to a single site binding model using MicroCal PEAQ-ITC analysis software, (d) Chart summarizing the *K_D_* values for all ATG8 swap chimera interactions tested, including two replicates with the PexRD54 AIM peptide and three replicates with full-length PexRD54.

### A single residue in the first β-strand underpins discriminatory binding to PexRD54

In parallel, to compare the AIM binding pockets of ATG8-2.2 and ATG8-4, we generated a homology model for ATG8-4-AIM peptide complex using our previous ATG8-2.2-AIM peptide complex as a template. ATG8-4 adopts a globular structure composed of C-terminal ubiquitin-like domain and N-terminal helical domain consisting of tandem α helices (α1 and α2) (Fig. 4a). Close inspection of ATG8-4-AIM peptide complex revealed that the AIM peptide is anchored in a cavity at the surface of ATG8-4 via (i) electrostatic interactions with ATG8-4 residues, and (ii) burial of AIM hydrophobic residues in two pockets of ATG8-4 (Fig. 4b-c). The Trp-378 of the AIM peptide sits in a hydrophobic pocket located between α2 and β1 of ATG8-4, whereas Val-381 resides in a distinct hydrophobic pocket between β2 and α3 (Fig. 4a-b; Fig. S11).

**Fig. 4.**
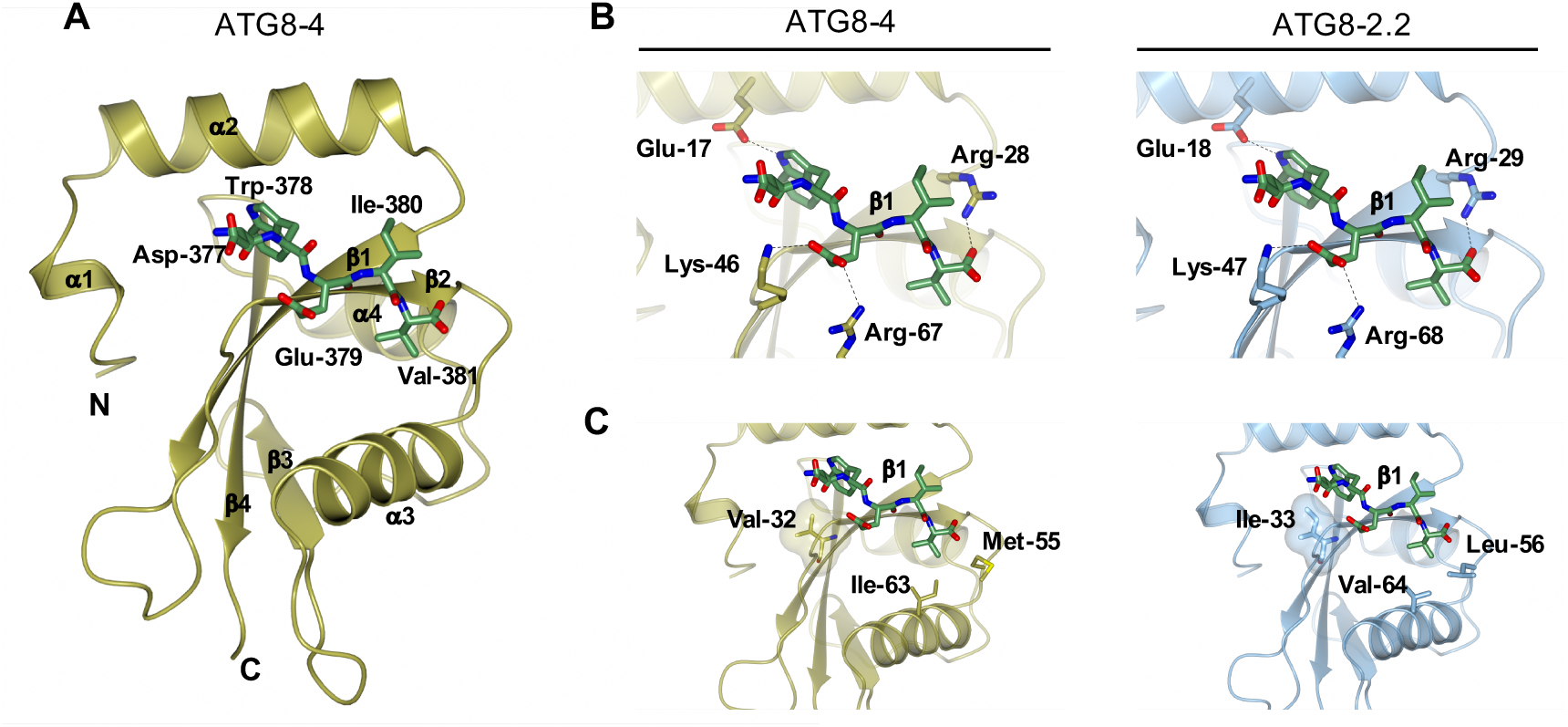
A Comparison of ATG8-2.2 structure and ATG8-4 model identifies polymorphic residues within the AIM binding site. (a) Homology model of ATG8-4 and PexRD54 AIM peptide complex. ATG8-4 and PexRD54 AIM are illustrated in cartoon and stick representation, α-helices, (β-strands, N‐ and C-termini of ATG8-4 are labelled, (b) Zoomed in view of the AIM peptide binding pocket of ATG8-4 (left) and ATG8-2.2 (right), with amino acids making electrostatic interactions labelled, (c) Zoomed in view of the AIM binding pocket of ATG8-4 (left) and ATG8-2.2 (right), highlighting differential residues contributing to hydrophobic interactions with the PexRD54 AIM peptide.

Comparison of the AIM binding pockets of ATG-2.2 and ATG8-4 revealed that amino acids mediating electrostatic interactions are conserved between both ATG8s (Fig. 4b). However, three residues that contribute to the hydrophobic pocket that accommodates the AIM peptide are polymorphic between ATG8-2.2 and ATG8-4 (Fig. 4c). Ile-33, which is located in the W pocket of ATG8-2.2, is changed to Val-32 in ATG8-4. Similarly, Leu-56 and Val-64, located in the Val-381 binding pocket of ATG8-2.2, are replaced by Met-55 and Ile-63, respectively. In sum, our structural analysis suggested that three polymorphic residues between ATG8-4 and ATG8-2.2 could contribute to the differential interactions with PexRD54 AIM.

To test if these residues underpin the binding specificity, we mutated each of these residues in the ATG8-4 background to match ATG8-2.2 (**Fig. 3a**). We also generated a combined triple mutant (ATG8-4-3x) and assayed all of these variants for gain-of-binding to PexRD54 (**Fig. 5a**). The ATG8-4 point mutant Val-32 to Ile-32 (ATG8-4-V32I), within the first β-strand, partially restored binding to PexRD54 in Co-IP experiments (**Fig. 5a**). We then purified ATG8-4-V32I variant and quantified the gain-of-binding phenotype using ITC (**Fig. S12a-b**). Remarkably, with both PexRD54 AIM peptide and the full-length PexRD54, the ATG8-4-V32I mutant showed a strong gain-of-binding phenotype, restoring binding to levels that are similar to ATG8-2.2 (**Fig. 5b-c**). We obtained similar results in multiple biological replicates (**Fig. S12c-e**). Altogether, these results suggest that a single amino acid residue, Val-32 in the first β-strand, determines differential binding affinity of ATG8-4 towards PexRD54.

**Fig. 5.**
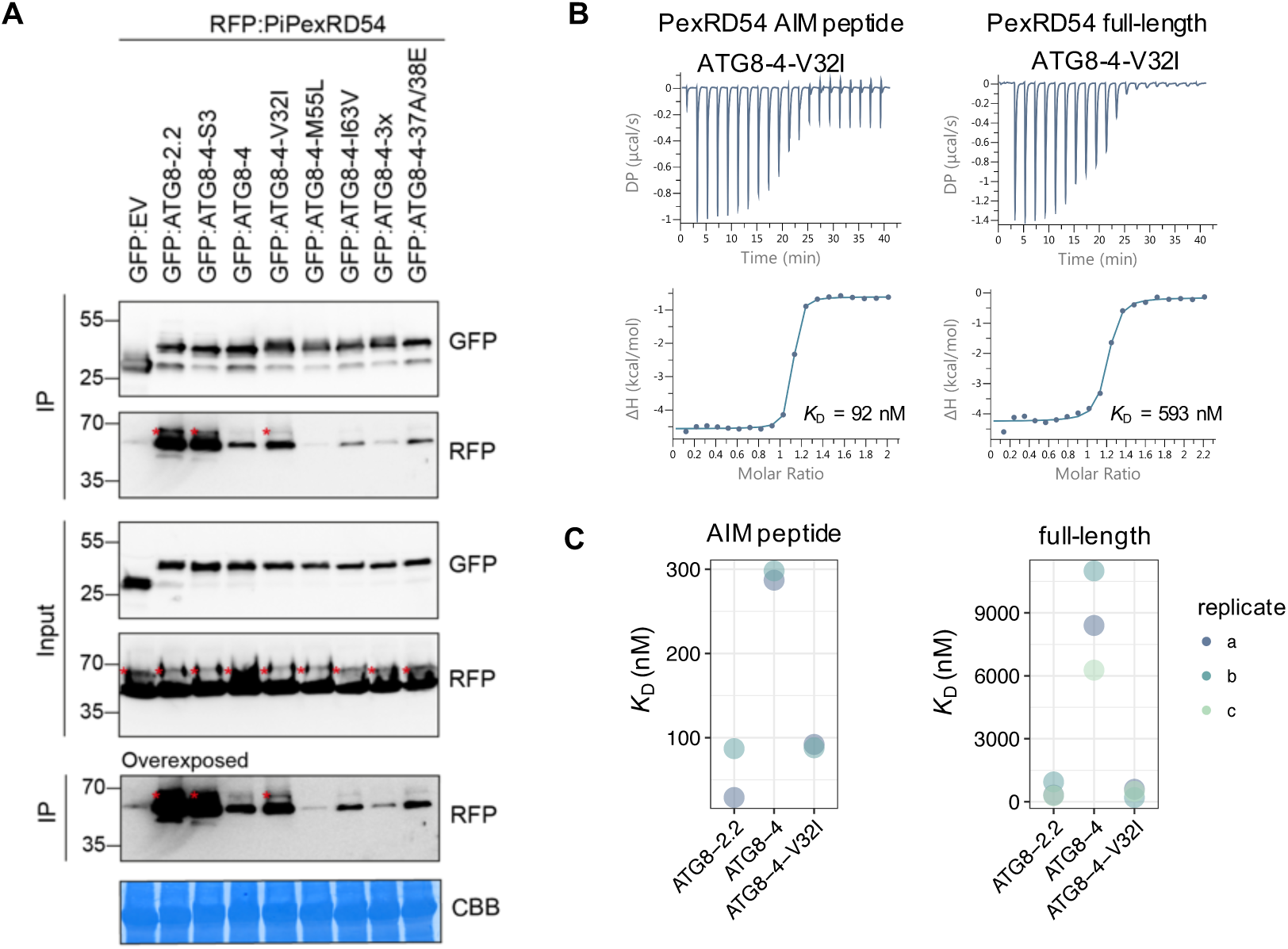
A single amino acid residue, Val-32 in the first β-strand, determines differential binding affinity of ATG8-4 towards PexRD54. (a) Co-immunoprecipitation experiment between PexRD54 and all ATG8-4 point mutants. RFP:PiPexRD54 was transiently co-expressed with the controls GFP:EV, GFP:ATG8-2.2, GFP:ATG8-4-S3, and GFP:ATG8-4, and all of the GFP:ATG8-4 point mutants. Immunoprecipitates (IPs) were obtained with anti-GFP antibody and total protein extracts were immunoblotted with appropriate antisera (listed on the right of each). Stars indicate expected band sizes. (b) The binding affinity of ATG8-4-V32I with PexRD54-AIM peptide and full-length PexRD54 was determined by ITC. The top panels show heat differences upon injection of ligands and lower panels show integrated heats of injection (•) and the best fit (solid line) to a single site binding model using MicroCal PEAQ-ITC analysis software. (c) Chart summarizing the *K_D_* values for each interaction, including two replicates with the PexRD54 AIM peptide and three replicates with full-length PexRD54.

### The N terminal β-strand defines the protein interactor profiles of ATG8-2.2 and ATG8-4

Since we found that the first β-strand contributes to selective binding to the AIM-containing substrate PexRD54, we hypothesized that this region also underpins binding specificity to other ATG8 interacting proteins. To test this, we performed *in planta* immunoprecipitation with tandem mass spectrometry (IP-MS) experiments with ATG8-4-S3, ATG8-2.2 and ATG8-4, resulting in a list of 291 proteins. Firstly, we detected significant overlap with our ATG8 interactome data (∼40% of interactors), validating our IP-MS approach (**Table S3**). We then interrogated this dataset to see if we could identify interactors enriched for interaction with either of the ATG8 isoforms. Similar to our first ATG8 interactome screen, ATG8-2.2 and ATG8-4 associated with distinct sets of proteins (**Fig. 6a**). Close to two-thirds of the proteins in the dataset were found to be significantly enriched in their interaction with either ATG8-2.2 (177 proteins) or ATG8-4 (6 proteins) using an ANOVA analysis with a post-hoc Tukey HSD, while remaining proteins similarly interacted with both isoforms (105 proteins) (**Table S4; Fig. 6c**). Much like the ATG8 interactome dataset, this dataset had an overrepresentation of predicted AIM-containing proteins (50%), with a majority of those AIMs evolutionarily conserved (70%), as compared to a random set of *N. benthamiana* proteins of the same size (35% and 27%, respectively) (**Fig. 6b**; **Table S4**; **Table S2**).

We then compared the interaction profile of ATG8-4-S3 to those of ATG8-2.2 and ATG8-4. The ATG8-4-S3 interaction profile more closely resembled ATG8-2.2 than ATG8-4, indicating that inclusion of the first β-strand from ATG8-2.2 in the ATG8-4 backbone was sufficient to shift the specificity of the resulting chimera (**Fig. 6a**). ATG8-4-S3 associated with around 40% of the proteins that were significantly enriched in the ATG8-2.2 pull-down to a level statistically indistinguishable from ATG8-2.2 (**Fig. 6d; Fig. S13a**). Within this set of ATG8-2.2 and ATG8-4-S3 enriched proteins, there was an overrepresentation of predicted AIM-containing proteins (53%), of which a majority of AIMs were conserved (83%), compared to the ATG8-2.2 enriched proteins that ATG8-4-S3 does not interact with (45% and 61%, respectively) (**Fig. 6d**). In addition, ATG8-4-S3 does not interact with two of the six ATG8-4 enriched interactors, and maintains similar interaction with all interactors common to both ATG8-2.2 and ATG8-4 (**Fig. S13b-c**). To see if the interactors significantly enriched in both the ATG8-2.2 and ATG8-4-S3 pull-downs had any specific properties, we performed gene ontology analyses, homology searches and localization predictions (**Table S4**). These analyses revealed that there are no other obvious trends within the group of proteins that seem to show preference for interaction with the ATG8-4-S3, besides the overrepresentation of proteins with evolutionarily conserved AIMs (**Table S4**).

**Fig. 6.**
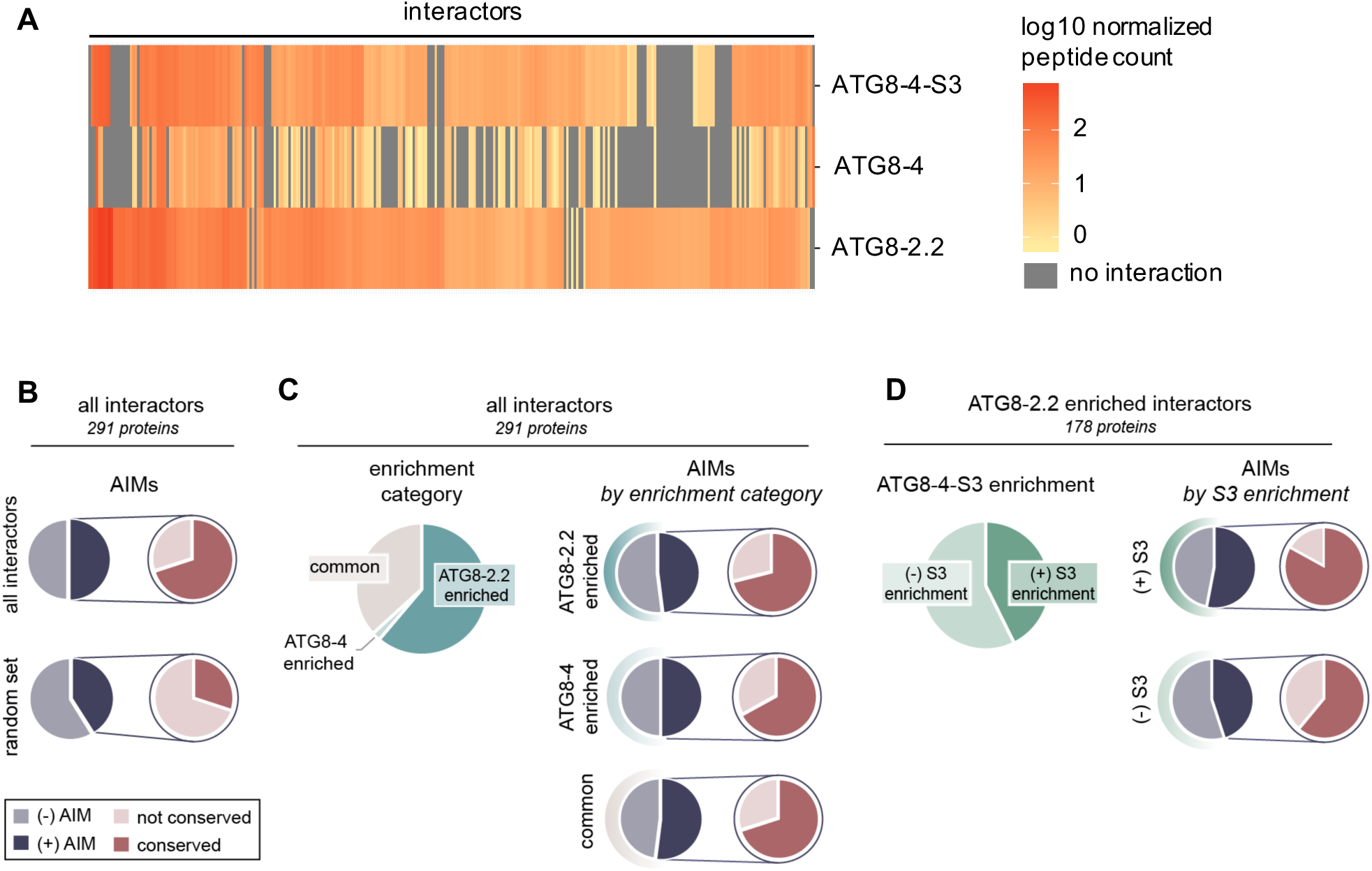
The first β-strand defines the AIM-dependent interaction profiles of ATG8 isoforms. (a) Heatmap showing the interaction profiles of ATG8-2.2, ATG8-4, and ATG8-4-S3. The average peptide count data was log10 normalized, and then used to construct a hierarchically clustered heatmap with the scale as shown, (b) All interactors inthedataset (291 proteins) and their closest *Arabidopsis thaliana* and *Marchantia polymorpha* homologs were analyzed for predicted AIMs using iLIR software [38]. The proportion of interactors that contain a predicted AIM—as well as the proportion of those AIMs that are conserved—are summarized. These are compared to the analogous values calculated from the average of three sets of random proteins from the *Nicotiana benthamiana* proteome (291 proteins/ set), (c) All interactors were divided into enrichment categories based on whether they showed a significantly (p<0.05) stronger interaction with ATG8-2.2 or ATG8-4 as determined by an ANOVA with a post-hoc Tukey’s test; interactors that showed no significant difference in their interaction with either protein were categorized as ‘common’. For each enrichment category, the proportion of interactors that contain a predicted AIM, and those AIMs that are conserved, are summarized, (d) For each interactor enriched in ATG8-2.2 pull-downs, we determined whether ATG8-4-S3 showed a significant (p<0.05) difference in its interaction strength compared to ATG8-2.2 using an ANOVA with a post-hoc Tukey’s test. Proteins that showed no statistical difference in their interaction with ATG8-4-S3 compared to ATG8-2.2 are categorized as ‘(+) S3 enrichment’; those that showed a statistical difference are categorized as ‘(-) S3 enrichment’. The proportion of interactors that contain a predicted AIM, and those AIMs that are conserved, are summarized for each ATG8-4-S3 enrichment category.

## Discussion

Over the last decade, our understanding of autophagy has evolved from a starvation-induced bulk degradation process to a highly selective cellular homeostasis pathway [6]. This begs the question— what are the molecular codes that define selective autophagy pathways? To date, the majority of the selective autophagy studies ascribed specificity to the cargo receptors that bind ATG8 via the conserved AIM [3]. In this study, we provide evidence that biochemical specialization of ATG8s is another layer of specificity that may contribute to functional specialization and subcellular compartmentalization of autophagy. This view is consistent with the evolutionary history of plant ATG8s, which have dramatically expanded in land plants, and exhibit family-specific clades that differ in fixed amino acid polymorphisms [32]. We also build on this view by producing an interactome resource for potato ATG8s that serves as a platform for further studies on the diversification of selective autophagy pathways.

As a part of our study, we generated a comprehensive ATG8 interactome resource for the six potato ATG8s. We validated the quality of our ATG8 interactome data in different ways. First, the results between the biological replicates were positively correlated, confirming reproducibility of our results (**Fig. S4**). We also noted that the ATG8 interactome contains most of the known ATG8 interacting proteins, such as core ATG proteins ATG4 and ATG7, and autophagy receptor NBR1 (**Table S1**). Moreover, four of the endogenous *N. benthamiana* ATG8s were present in the interactome (**Table S1**). One of the ATG8s, NbATG8-4, exhibited a selective interaction pattern, interacting almost exclusively with ATG8-4, its closest homolog (**Fig. S8**). This supports our model that autophagosomes likely carry different populations of ATG8s. The interactome also has several vesicle trafficking components, such as Rab GTPase activating proteins and coatomer subunits, which were also uncovered in human ATG8 interactome studies [22,23]. The dataset also contains highly abundant proteins such as ribosome and proteasome subunits, which are known to undergo autophagic degradation [39,40]. However, further studies will be necessary to distinguish these particular interactors from the CRAPome, the common false positive proteins observed in immunoprecipitation experiments [41].

Interestingly, we detected several proteins that have not yet been associated with autophagy. Notable interactors include dual specificity protein tyrosine kinase, web family, and lipin family proteins (**Table S1**). As these proteins contain conserved predicted AIMs, our interactome could serve as a great resource for future studies that aim to discover novel autophagy receptors, adaptors, and regulators. Relatedly, the overrepresentation of interactors in the dataset that contain predicted evolutionarily conserved AIMs suggests that the interactome is enriched in putative autophagic components and indicates that evolutionary conservation may be an additional parameter to predict *bona fide* AIM sequences.

Recent studies indicate that human ATG8s are functionally specialized. The interactor profiles of human ATG8 isoforms has been explored using IP-MS [22,23] and proximity labeling [28] proteomics. These studies revealed limited overlap between different ATG8 isoforms, particularly with the proximity labeling proteomics, which primarily identifies cargoes within various ATG8 autophagosomes [28]. In addition, mechanistic studies on ATG8 interacting proteins also support specialization of human ATG8 isoforms. For example, the xenophagy receptor NDP52 specifically binds the ATG8 protein LC3C [42], whereas the autophagy adaptor PLEKHM1 prefers to bind a different ATG8, GABARAP [16]. A recent study even revealed GABARAP interaction motif (GIM) that mediates high affinity binding of GABARAP-interacting proteins [25]. Considering that some plant species have over 20 ATG8 isoforms, compared to 6 in humans, it is reasonable to hypothesize that ATG8s directly contribute to functional diversification of selective autophagy pathways in plants. Indeed, our finding that ATG8s differentially interact with different types of cargo is consistent with this view (**Fig. 1c**; **Fig. S7**).

Previous studies investigating ATG8-AIM peptide complexes have shown that hydrophobic and electrostatic interactions underlie substrate specificity of ATG8 isoforms [24,29,43]. For example, a recent study that structurally characterized the *C. elegans* ATG8 isoforms LGG1 and LGG2 showed that the hydrophobic AIM binding pocket and the surrounding regions determine substrate specificity [24]. In contrast, since all the residues that form electrostatic interactions are conserved between ATG82.2 and ATG8-4, our data suggest hydrophobic interactions drive the marked difference in interaction strength between ATG8-2.2 and ATG8-4 with the substrate PexRD54. Future studies will reveal whether electrostatic interactions also contribute to substrate specificity of other plant ATG8 isoforms.

Detailed analysis of the ATG8-2.2 hydrophobic pocket revealed that isoleucine 33 (Ile-33)—which is only a methyl group larger than the valine residue found in the ATG8-4 isoform—primarily underlies binding to PexRD54, especially in *in vitro a*ssays. The isoleucine in ATG8-2.2 may better fill the hydrophobic cavity, and thus lead to stronger interactions with the PexRD54 AIM peptide. Ile-33 is highly conserved in various ATG8 isoforms [32], suggesting that the valine polymorphism (Val-32) in ATG8-4 is a recently derived polymorphism. It is tempting to speculate that this polymorphism was selected to evade PexRD54 binding, henceforth subversion of autophagy by the pathogen. Complementation of higher order ATG8 mutants with valine substituted ATG8 isoforms could challenge this hypothesis, and reveal whether valine substitution has an adverse effect on autophagy in general. Interestingly, a similar isoleucine to valine polymorphism underlies the pyrabactin selectivity of the ABA receptors PYL1 and PYL2 [44], highlighting how subtle differences in substrate binding pockets could lead to functional diversity.

The ATG8 N-terminal β-strand also underpins binding specificity to plant substrates, with polymorphisms in this region accounting for differential interaction with about 80 plant proteins. Proteins significantly enriched in both the ATG8-2.2 and ATG8-4-S3 pull-downs, compared to ATG8-4, had a higher proportion of predicted conserved AIMs (44%) than proteins that were enriched in ATG8-2.2, but not ATG8-4-S3 (27%). This suggests the N-terminal β-strand mediates discriminatory binding to AIM-containing substrates and leaves the door open for future studies to look at the structural basis of ATG8 substrate specificity determined via other interfaces.

In summary, plant ATG8 isoforms are specialized and bind distinct sets of proteins, and the hydrophobic pocket that accommodates the AIM peptide contributes to binding specificities. This suggests that multiple ATG8 isoforms should be assessed when measuring autophagy dynamics in plants. Based on this work, and other studies from mammalian systems, we propose a model where, together with the autophagy receptors, different ATG8 isoforms could contribute to the subcellular compartmentalization of various selective autophagy pathways, especially when they are active at the same time in a cell.

## Materials and Methods

### Gene cloning

#### Cloning for recombinant protein production for in vitro studies

PexRD54 was cloned in a previous study [33]. DNA encoding all different members of ATG8 family from *Solanum tuberosum,* amino acid residues Ser5-Asn114, (except for ATG8-4, Thr4-Lys113) were amplified by PCR from cDNA. For gain-of-function swaps in the ATG8-4 background, amino acid residues Thr4-Lys113, (except for ATG8-4-Swap1, Ser5-Lys113) were amplified as described above. All PCR amplicons were subsequently cloned into the pOPINE vector [45] using In-Fusion cloning based on in vitro homologous recombination, using a commercial kit (Clontech, Mountain View, CA, United States) generating uncleavable C-terminal 6xHis-tagged proteins for purification. ATG8-4-V32I mutant was custom synthesized into the pUC57-Amp vector (Genewiz, UK), and subsequently amplified and cloned into the pOPINE vector as described above. All constructs in pOPINE vectors, pOPINE-ATG8-2.2, pOPINE-ATG8-1.1, pOPINE-ATG8-1.2, pOPINE-ATG8-2.1, pOPINE-ATG8-3.1, pOPINE-ATG8-3.2, pOPINE-ATG8-4, pOPINE-ATG8-2.2, pOPINE-ATG8-4-Swap1, pOPINE-ATG8-4-Swap2, pOPINE-ATG8-4-Swap3, pOPINE-ATG8-4-Swap7 and ATG8-4-V32I, were transformed into *E. coli* strain BL21 (DE3) for recombinant protein production. All primers used in PCR for cloning are shown in **Table S5**. All the constructs were verified by DNA sequencing.

#### Cloning for in planta Co-Immunoprecipitations

The ATG8 isoforms were amplified from *Solanum tuberosum* cDNA and cloned into pK7WG2 vectors as described previously for ATG82.2 [33]. ATG8-2.2 and ATG8-4 were also cloned as level 0 modules for Golden Gate cloning [46]. ATG8 swaps and ATG8-4 point mutants were synthesized as level 0 modules for Golden Gate cloning [46]. GFP:ATG8-2.2, GFP:ATG8-4, GFP:ATG8 swaps and GFP:ATG8-4 point mutants were generated by Golden Gate assembly with pICSL13001 (long 35s promoter, The Sainsbury Laboratory (TSL) SynBio), pICSL30006 (GFP, TSL SynBio), pICH41414 (35s terminator, TSL SynBio), into the binary vector pICH47732 [46]. All constructs were verified by DNA sequencing.

#### Transient protein expression in N. benthamiana and total protein isolation

Transient gene expression *in planta* was performed by delivering T-DNA constructs with *Agrobacterium tumefaciens* GV3101 strain into 3- 4 week-old *N. benthamiana* plants as described previously [47]. *A. tumefaciens* strains carrying the plant expression constructs were diluted in agroinfiltration medium [10 mM MgCl_2_, 5 mM 2-(N-morpholine)-ethanesulfonic acid (MES), pH 5.6] to a final OD_600_ of 0.2, unless stated otherwise. For transient co-expression assays, *A. tumefaciens* strains were mixed in a 1:1 ratio. *N. benthamiana* leaves were harvested 3 days after infiltration, and protein isolation was conducted as previously described [47].

#### Heterologous protein production and purification for in vitro experiments

Bacteria expressing heterologous proteins were grown in auto-induction media (AIM) at 37 °C and transferred to 16 °C overnight upon induction. Cell pellets were resuspended in buffer A1 (50 mM Tris-HCl pH 8, 500 mM NaCl, 50 mM glycine, 5% (v/v) glycerol, 20 mM imidazole, and EDTA free protease inhibitor). The cells were lysed by sonication and subsequently spun to produce the clear lysate. A single Ni2+-NTA capture step was followed by gel filtration onto a Superdex 75 26/600 gel filtration column pre-equilibrated in buffer A4 (20 mM HEPES pH 7.5, 150 mM NaCl). The fractions containing His tagged ATG8 protein of interest were pooled and concentrated as appropriate, and the final concentration was judged by absorbance at 280 nm (using a calculated molar extinction coefficient of each protein). The purity of proteins was judged by running 16% SDS-PAGE gels and stained with instant blue. PexRD54 was purified as described previously [34].

### Protein-protein interaction studies

#### Isothermal Titration Calorimetry

All calorimetry experiments were recorded using a MicroCal PEAQ-ITC (Malvern, UK). To test the interaction of ATG8 proteins with full length PexRD54, experiments were carried out at 18 °C using 20 mM HEPES pH 7.5, 500mM NaCl buffer. The calorimetric cell was filled with 110 μM PexRD54 and titrated with 1.1 mM ATG8 protein. For the PexRD54 AIM peptide studies, all experiments were conducted at 25 °C using 20 mM HEPES pH 7.5, 150mM NaCl buffer. The calorimetric cell was filled with 110 μM of ATG8 titrated with 1.1 mM of peptide. For each ITC run, a single injection of 0.5 μl of ligand was followed by 19 injections of 2 μl each. Injections were made at 120 s intervals with a stirring speed of 750 rpm. The raw titration data were integrated and fitted to a one-site binding model using the built-in software of MicroCal PEAQ ITC.

#### Co-immunoprecipitation and immunoblot analyses

The Co-IP protocol described by Win et al. 2011 was adapted for GFP and RFP fusions with the following modifications. Immunoprecipitation was performed by affinity chromatography with GFP_Trap_A beads (Chromotek), and elution of the proteins from the beads was performed by heating 10 minutes at 70°C. Proteins were separated by SDS-PAGE and were transferred onto a polyvinylidene diflouride membrane using a Trans-Blot turbo transfer system (Bio-Rad, Munich). The membrane was blocked with 5% milk in Tris-buffered saline and Tween 20. GFP detection was performed in a single step by a GFP (B2):sc-9996 horseradish peroxidase (HRP)-conjugated antibody (Santa Cruz Biotechnology, Santa Cruz, CA, U.S.A.); RFP detection was performed with a rat anti-RFP 5F8 antibody (Chromotek, Munich) and a HRP-conjugated anti-rat antibody. Membrane imaging was carried out with an ImageQuant LAS 4000 luminescent imager (GE Healthcare Life Sciences, Piscataway, NJ, U.S.A.). Instant Blue (Expedeon, Cambridge) staining of the rubisco was used as a loading control.

### Mass spectrometry

#### (i) ATG8 interactome

##### Sample preparation for mass spectrometry

Following the protein purification and washing steps, the beads were resuspended in 2 bead volumes of 100 mM ammonium bicarbonate (Fluka 09830-500G). Proteins were digested from the beads by addition of 400 ng Lys-C (Wako PEF 7041) and incubation on a Thermoshaker with 1300 rpm for 4 hours at 37°C. Subsequently, the supernatant was transferred to a fresh tube and reduced in 0.6 mM TCEP-HCl (Tris 2-carboxyethyl phosphine hydrochloride, Sigma 646547-10 x 1ml) for 30 min at 60°C followed by an alkylation reaction in 4 mM MMTS (methyl methanethiosulfonate, Fluka 64306) for 30 min at room temperature in the dark. Peptides were further digested by addition of 400 ng Trypsin (Trypsin Gold Promega V5280) and overnight incubation at 37°C. The digest was stopped by addition of TFA (trifluoroacetic acid, Aldrich T63002) to a final concentration of 1%.

##### NanoLC-MS Analysis

The nano HPLC system used was an UltiMate 3000 RSLC nano system (Thermo Fisher Scientific, Amsterdam, Netherlands) coupled to a Q Exactive HF-X mass spectrometer (Thermo Fisher Scientific, Bremen, Germany), equipped with a Proxeon nanospray source (Thermo Fisher Scientific, Odense, Denmark). Peptides were loaded onto a trap column (Thermo Fisher Scientific, Amsterdam, Netherlands, PepMap C18, 5 mm × 300 μm ID, 5 μm particles, 100 Å pore size) at a flow rate of 25 μL min-1 using 0.1% TFA as mobile phase. After 10 min, the trap column was switched in line with the analytical column (Thermo Fisher Scientific, Amsterdam, Netherlands, PepMap C18, 500 mm × 75 μm ID, 2 μm, 100 Å). Peptides were eluted using a flow rate of 230 nl min-1, and a binary 3h gradient, respectively 225 min.

The gradient starts with the mobile phases: 98% A (water/formic acid, 99.9/0.1, v/v) and 2% B (water/acetonitrile/formic acid, 19.92/80/0.08, v/v/v), increases to 35%B over the next 180 min, followed by a gradient in 5 min to 90%B, stays there for 5 min and decreases in 2 min back to the gradient 98%A and 2%B for equilibration at 30°C.

The Q Exactive HF-X mass spectrometer was operated in data-dependent mode, using a full scan (m/z range 350-1500, nominal resolution of 60,000, target value 1E6) followed by MS/MS scans of the 10 most abundant ions. MS/MS spectra were acquired using normalized collision energy of 28, isolation width of 1.0 m/z, resolution of 30.000 and the target value was set to 1E5. Precursor ions selected for fragmentation (exclude charge state 1, 7, 8, >8) were put on a dynamic exclusion list for 60 s. Additionally, the minimum AGC target was set to 5E3 and intensity threshold was calculated to be 4.8E4. The peptide match feature was set to preferred and the exclude isotopes feature was enabled.

##### Data Processing and peptide identification

For peptide identification, the RAW-files were loaded into Proteome Discoverer (version 2.1.0.81, Thermo Scientific). All hereby created MS/MS spectra were searched using MSAmanda v2.1.5.9849, Engine version v2.0.0.9849 [48]. For the 1st step search the RAW-files were searched against a *N. benthamiana* genome database called Nicotiana_Benthamiana_Nbv6trPAplusSGNUniq_20170808 (398,682 sequences; 137,880,484 residues), supplemented with common contaminants, using the following search parameters: The peptide mass tolerance was set to ±5 ppm and the fragment mass tolerance to 15ppm. The maximal number of missed cleavages was set to 2, using tryptic enzymatic specificity. The result was filtered to 1 % FDR on protein level using Percolator algorithm integrated in Thermo Proteome Discoverer. A sub-database was generated for further processing.

For the 2nd step the RAW-files were searched against the created sub-database (36,152 sequences; 16,892,506 residues), using the following search parameters: Beta-methylthiolation on cysteine was set as a fixed modification, oxidation on methionine, deamidation on asparagine and glutamine, acetylation on lysine, phosphorylation on serine, threonine and tyrosine, methylation and di-methylation on lysine and arginine, tri-methylation on lysine, ubiquitination on lysine were set as variable modifications. Monoisotopic masses were searched within unrestricted protein masses for tryptic enzymatic specificity. The peptide mass tolerance was set to ±5 ppm and the fragment mass tolerance to ±15 ppm. The maximal number of missed cleavages was set to 2. The result was filtered to 1% FDR on peptide level using Percolator algorithm integrated in Thermo Proteome Discoverer. The localization of the post-translational modification sites within the peptides was performed with the tool ptmRS, based on the tool phosphoRS [49]. Peptide areas have been quantified using in-house-developed tool APQuant. The mass spectrometry proteomics data have been deposited to the ProteomeXchange Consortium via the PRIDE [50, 51] partner repository with the dataset identifier PXD011226.

##### Data filtering

For each assayed construct the PSM values were averaged between replicates, and then the dataset was filtered to remove any proteins that showed PSM value of >10 with the empty vector control, as well as any proteins where none of the ATG8-GFP fusions exhibited an average PSM value >10. After this basic filtering, the dataset was further filtered such that for all interactors with an EV average PSM value >4, at least one of the ATG8-GFP fusions had to exhibit >3x the EV PSM value (e.g. for EV average PSM = 9.5, one ATG8 > 28.5). This resulted in a final list of 621 proteins (**Table S1**).

##### Network analysis

For each interactor in the dataset, the closest *A. thaliana* homolog was predicted using BLAST, and the gene ontology (GO) annotations were obtained using Blast2GO [52]. The GO annotations were used to reduce the complexity of the final interactome presented in Table S1. The interactors were sorted by cellular compartment or biological process, respectively, and then collapsed based on shared annotations. The number of interactors in each shared annotation group was recorded, and the average PSM values were calculated for each group for every ATG8; the resulting tables were imported into Cytoscape [53] to make the network figures. The former values were used to scale the node sizes across all network representations, and the latter values were used to weight the edges for each individual ATG8-group connection.

#### (ii) ATG8-4-S3 mutant analysis

##### Sample preparation for mass spectrometry

Immunoprecipitated protein samples were separated by SDS-PAGE (4-20% gradient gel, Biorad), and after staining with Coomassie brilliant Blue G-250 CBB (SimplyBlue Safe stain, Invitrogen) the proteins were cut out and gel slices were destained in 50% acetonitrile. Reduction and alkylation was done by incubation for 45 min in 10 mM DTT, followed for 30 min in the dark in 55 mM chloroacetamide. After several washes with 25 mM ammonium bicarbonate, 50% acetonitrile gel slices were dehydrated in 100% acetonitrile. Gel pieces were rehydrated with 50 mM ammonium bicarbonate and 5% acetonitrile containing 20 ng/μl trypsin (Pierce), and digestion proceeded overnight at 37 °C. Tryptic peptides were sonicated from the gel in 5% formic acid, 50% acetonitrile, and the total extracts were evaporated until dry.

##### Data processing and peptide identification

IP-MS analysis of the GFP control and ATG8-GFP IP samples was done as previously described [54]. Briefly, LC-MS/MS analysis was performed with a Orbitrap Fusion Trihybrid mass spectrometer (Thermo Scientific) and a nanoflow-HPLC system (Dionex Ultimate3000, Thermo Scientific) described previously [54]. The peptide identification was performed by searching the in-house *N. benthamiana* database Nicotiana_Benthamiana_Nbv6trPAplusSGNUniq_20170808 (398,682 sequences; 137,880,484 residues) using Mascot (v 2.4.1 Matrix Science) with the modification of allowing Trypsin peptide termini. Scaffold (v4; Proteome Software) was used to validate MS/MS-based peptide and protein identifications and annotate spectra using a search criteria of a minimum of two peptides with MASCOT ion scores above 20 and 95% peptide identity. Selected spectra were manually inspected before acceptance. The mass spectrometry proteomics data have been deposited to the ProteomeXchange Consortium via the PRIDE [50, 51] partner repository with the submission reference 1-20181024-42155 and the formal identifier to be obtained upon dataset acceptance.

##### Data filtering

The peptide count values were first normalized to the peptide counts for GFP in each sample to adjust for varying expression levels. Then, after averaging the replicate values for each assayed construct, the dataset was filtered to remove any proteins that showed a peptide count of >6 with the empty vector control, as well as any proteins where none of the ATG8-GFP fusions exhibited a peptide count >10. This resulted in a list of 496 proteins. This dataset was further filtered by removing proteins that showed extreme unevenness among replicates, resulting in a final ATG8-4-S3 dataset of 291 proteins amenable to statistical analysis (**Fig. S14**).

### Protein confirmation

#### Intact mass spectrometry for accurate mass determination

LC-MS analysis, performed using standard procedures within the John Innes Centre proteomics facility, confirmed that each of the heterologously expressed proteins had the molecular mass as expected from the expressed sequence (**Fig. S9b, S10b, S12b**).

#### Structural homology modelling

Due to high sequence identity, ATG8-2.2 was used a template to generate a homology model of ATG8-4. The amino acid sequence of ATG8-4 was submitted to Protein Homology Recognition Engine V2.0 (Phyre^2^) for modelling [55]. The coordinates of ATG8-2.2 structure (5L83) were retrieved from the Protein Data Bank (PDB) and assigned as modelling template by using Phyre^2^ Expert Mode. The resulting model of ATG8-4 comprised amino acids Thr-4 to Glu-112 and was illustrated in CCP4MG software [56].

## Acknowledgements

This work was supported by the European Research Council Advanced Investigator Grant NGRB, Biotechnology and Biological Sciences Research Council grants BB/J004553 and BB/P012574, The Gatsby Charitable Foundation, John Innes Foundation, Austrian Academy of Sciences, WWTF (Project No: LS17-047), Austrian Science Fund (SFB F3402, TRP 308-N15), and the international Project ERA-CPAS (I 3686). We acknowledge Joe Win for assembling the *N. benthamiana* database and Peter Waterhouse for making the V6 annotation of the *N. benthamiana* transcriptome available prior to publication.

---

**Fig. S1.**
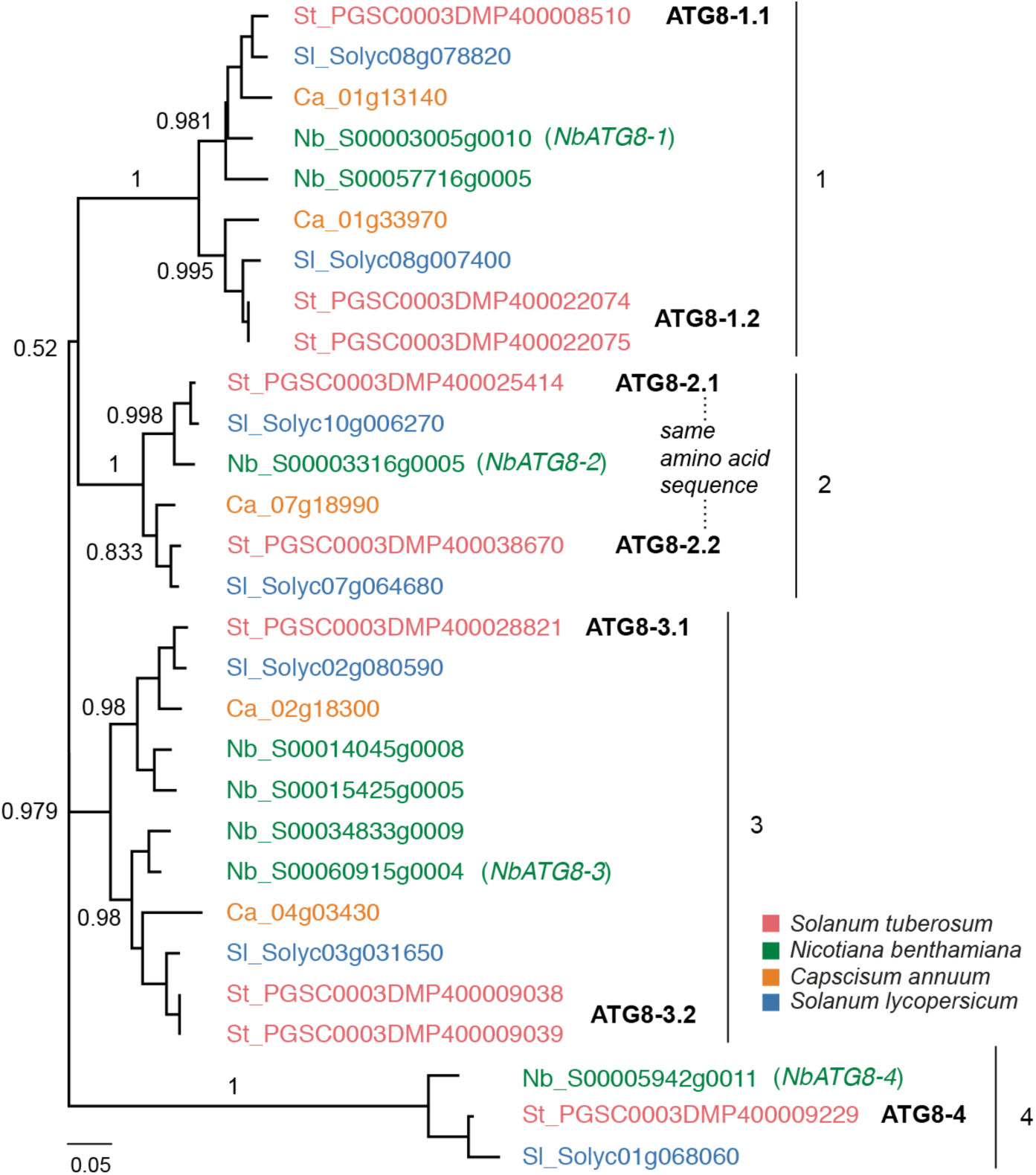
Orthologous relationships between Solanaceous ATG8 isoforms. A more detailed view of Fig. 1a. Unrooted maximum-likelihood phylogenetic tree of 29 ATG8 homologs with clades marked on the right, and colors indicating plant species. The tree was calculated in MEGA7 [36] from a 369 nucleotide alignment (MUSCLE [37], codon-based). The bootstrap supports of the major nodes are indicated. The scale bar indicates the evolutionary distance based on nucleotide substitution rate.

**Fig. S2.**
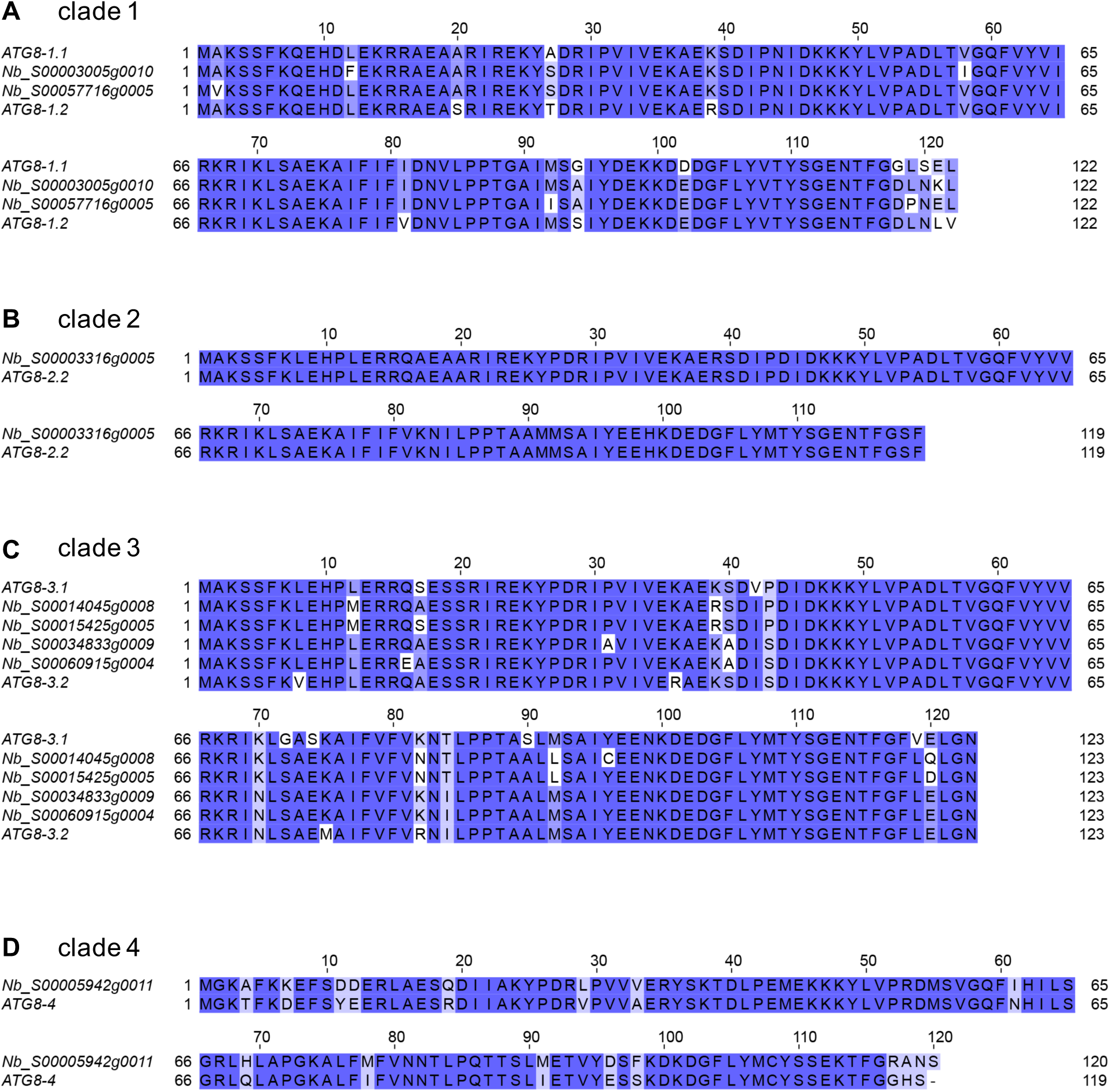
Orthologous relationships between *Solanum tuberosum* and *Nicotiana benthamiana* ATG8 isoforms. (a-d) Alignments of *S. tuberosum* and *N. benthamiana* ATG8s by clade (MUSCLE [37]), visualized with Jalview. *S. tuberosom* ATG8s are named as in **Fig. S1**; (b) only the *S. tuberosum* ATG8-2.2 is shown for the clade 2 alignment, as both ATG8-2.1 and ATG8-2.2 have the same amino acid sequence.

**Fig. S3.**
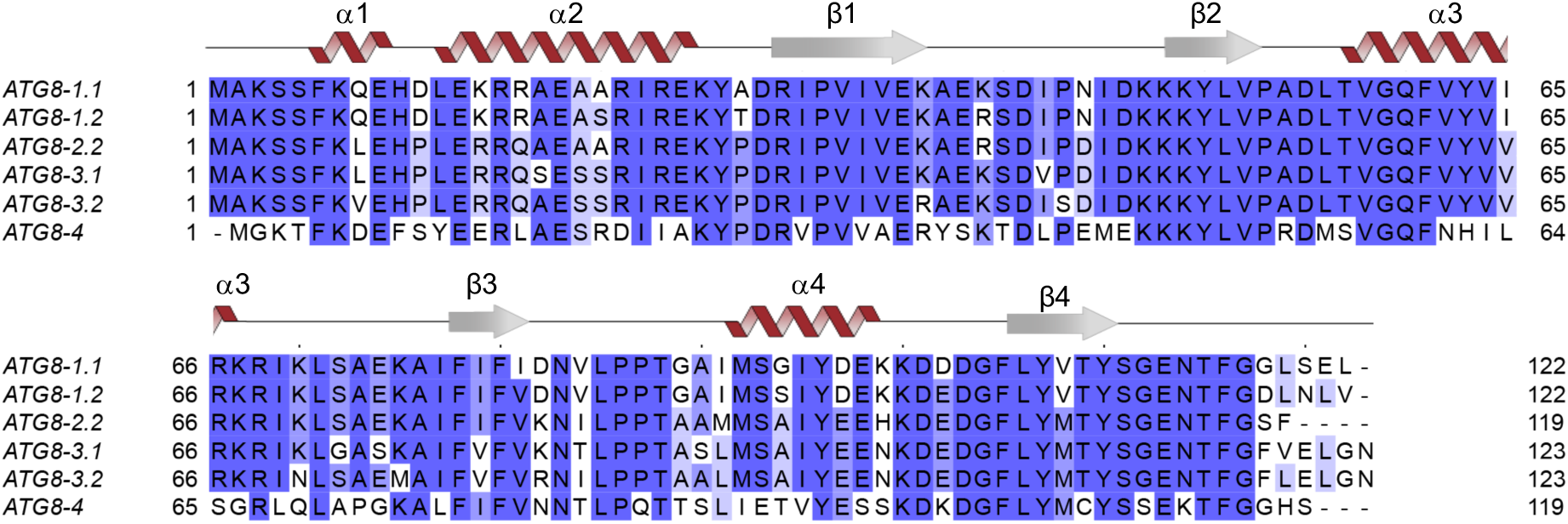
Sequence diversity among potato ATG8 isoforms. Alignment of all *S. tuberosum* ATG8s (MUSCLE [37]), visualized with Jalview, with the protein model above corresponding to the ATG8-2.2 structure. ATG8s are named as in Fig. S1; only ATG8-2.2 is included in the alignment, as both ATG8-2.1 and ATG8-2.2 have the same amino acid sequence.

**Fig. S4.**
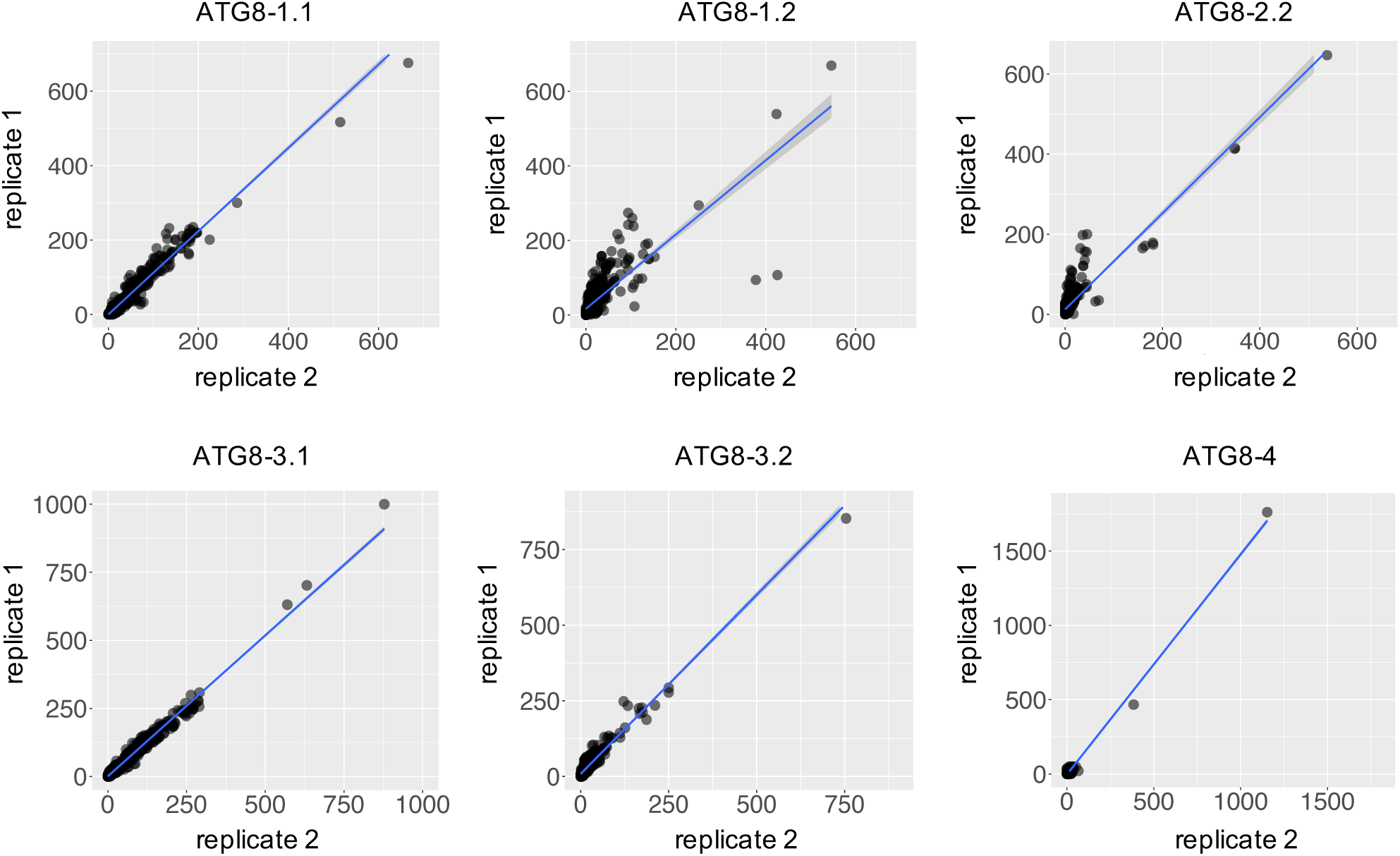
ATG8 interactome data is reproducible across replicates. The PSM values for the two replicates of each ATG8 isoform in the interactome dataset (621 interactors) were plotted in a pairwise fashion with a line of best fit, showing reproducibility across the replicates.

**Fig. S5.**
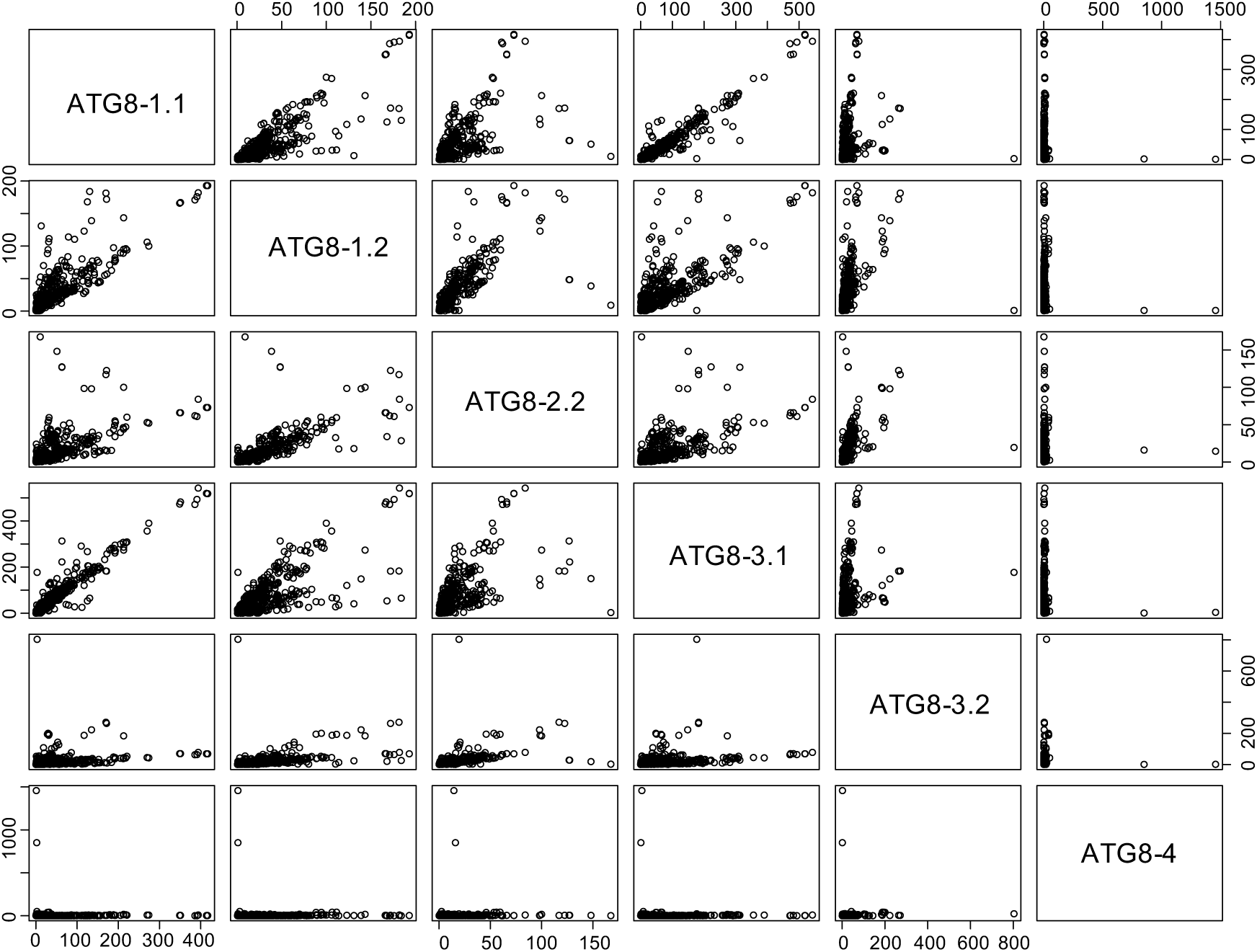
Solanaceous ATG8 isoforms have distinct protein interaction profiles. The average PSM values for each ATG8 isoform in the interactome dataset (621 interactors) were used to generate a correlation matrix, showing distinct interaction profiles for each ATG8, with varying degrees of overlap.

**Fig. S6.**
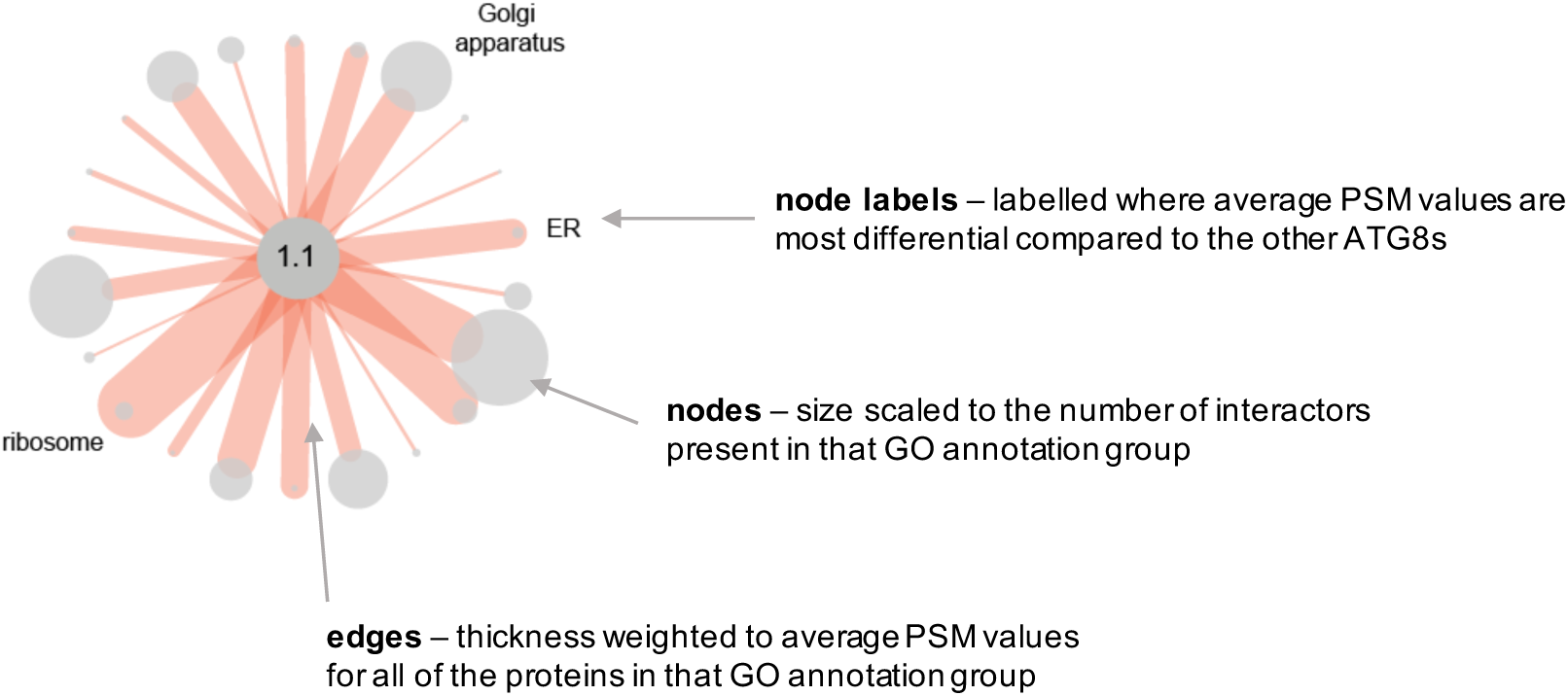
Graphical abstract for ATG8 interactome network representations. For both Fig. 1c and Fig. S7, nodes are scaled to the number of interactors present in each respective gene ontology (GO) annotation group, and edges are weighted to the average PSM values for all of the proteins in that GO annotation group for each ATG8. Nodes are labelled where the average PSM values are most differential when compared to other ATG8s.

**Fig. S7.**
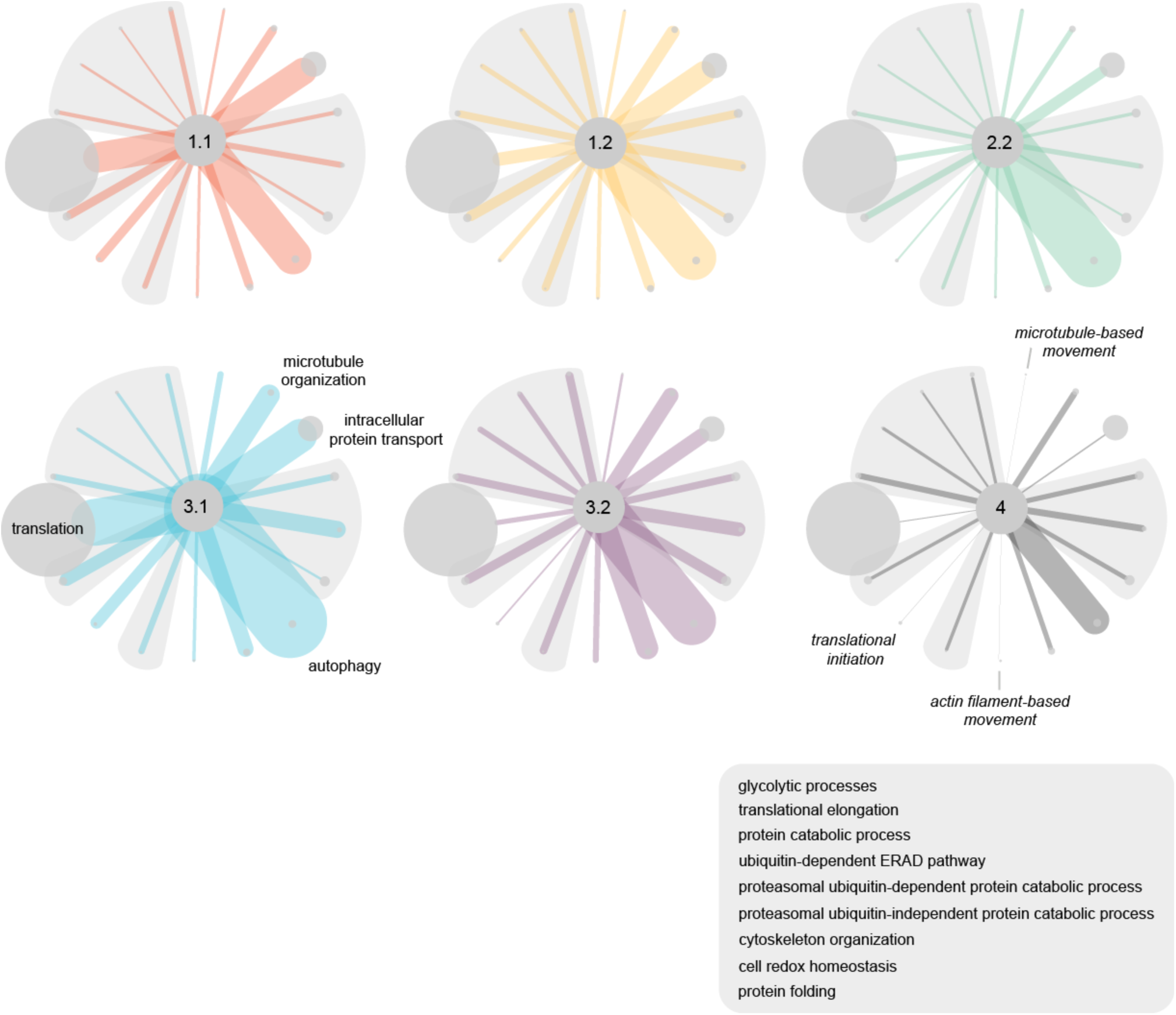
Network representation of the interactions between ATG8s and protein groups defined by biological process gene ontology (GO) annotations. For each interactor in the dataset, the closest *A. thaliana* homolog was predicted using BLAST, and the gene ontology annotations were obtained using Blast2GO [52]. Proteins were grouped based on the cellular compartment terms, and a subset of groups were chosen for representation. The sizes of the nodes are scaled to the number of interactors in each respective group, and the edges are weighed to the average PSM values for all the interactors in each respective group for each ATG8. Nodes are labelled where the average PSM value is most differential compared to the other ATG8s. Nodes shaded in gray exhibit similar average PSM values between all ATG8s, and the labels for these are included in the gray box. Figure S6 provides a graphical figure legend.

**Fig. S8.**
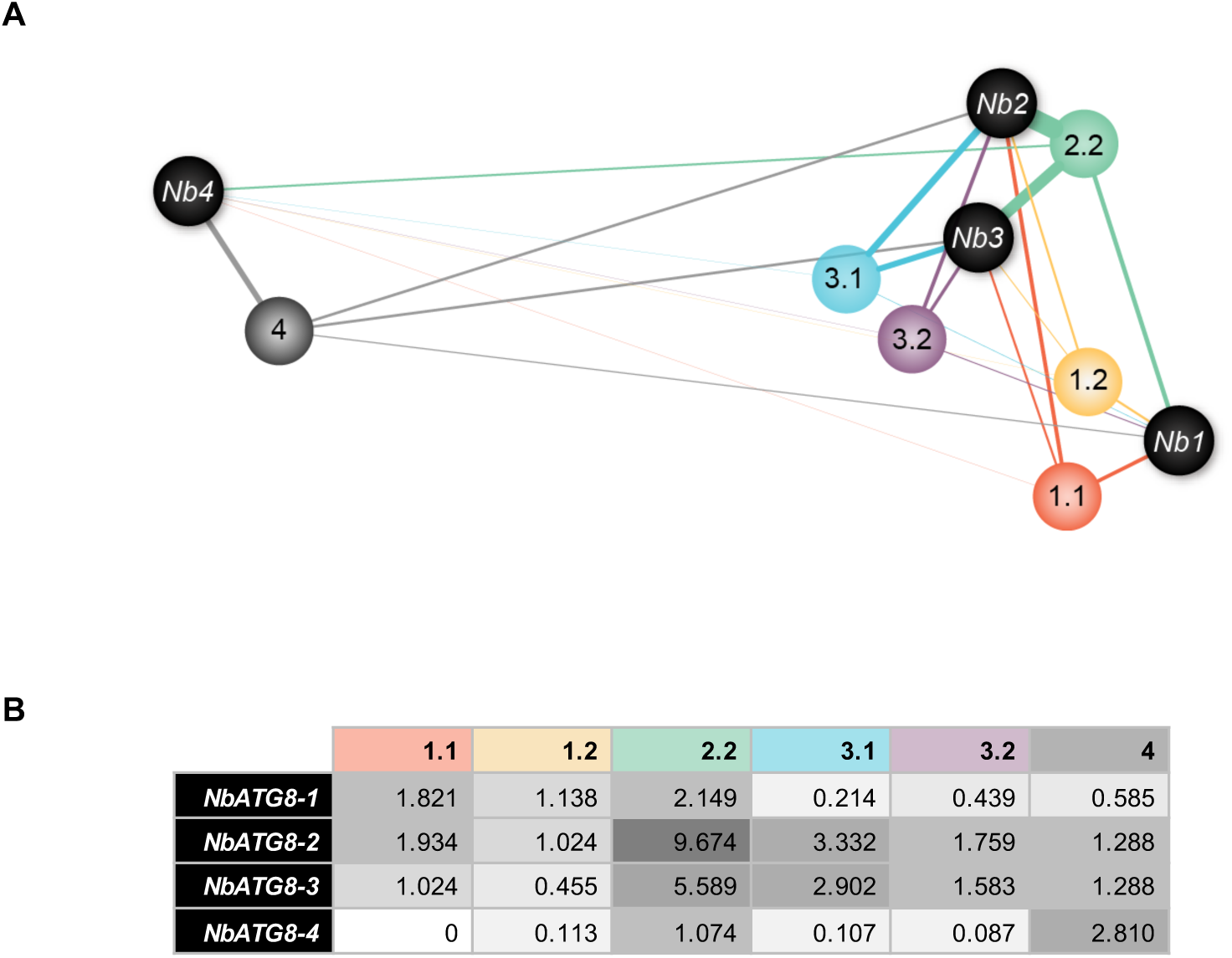
Network representation of interaction between potato ATG8s and endogenous *N. benthamiana* ATG8s. (a) Network representation of the interactions between potato ATG8s and endogenous *N. benthamiana* ATG8s. The edge widths are weighted to the GFP normalized peptide counts shown in (b). The spatial relationships between the ATG8s are approximately scaled to amino acid sequence identity, with more sequence related ATG8s clustering together, using Cytoscape [53]. The four *N. benthamiana* ATG8s present in the ATG8 interactome dataset—labelled here NbATG8-1- NbATG-4—are correspondingly labelled in Table S1 and Fig. S1 for reference.

**Fig. S9.**
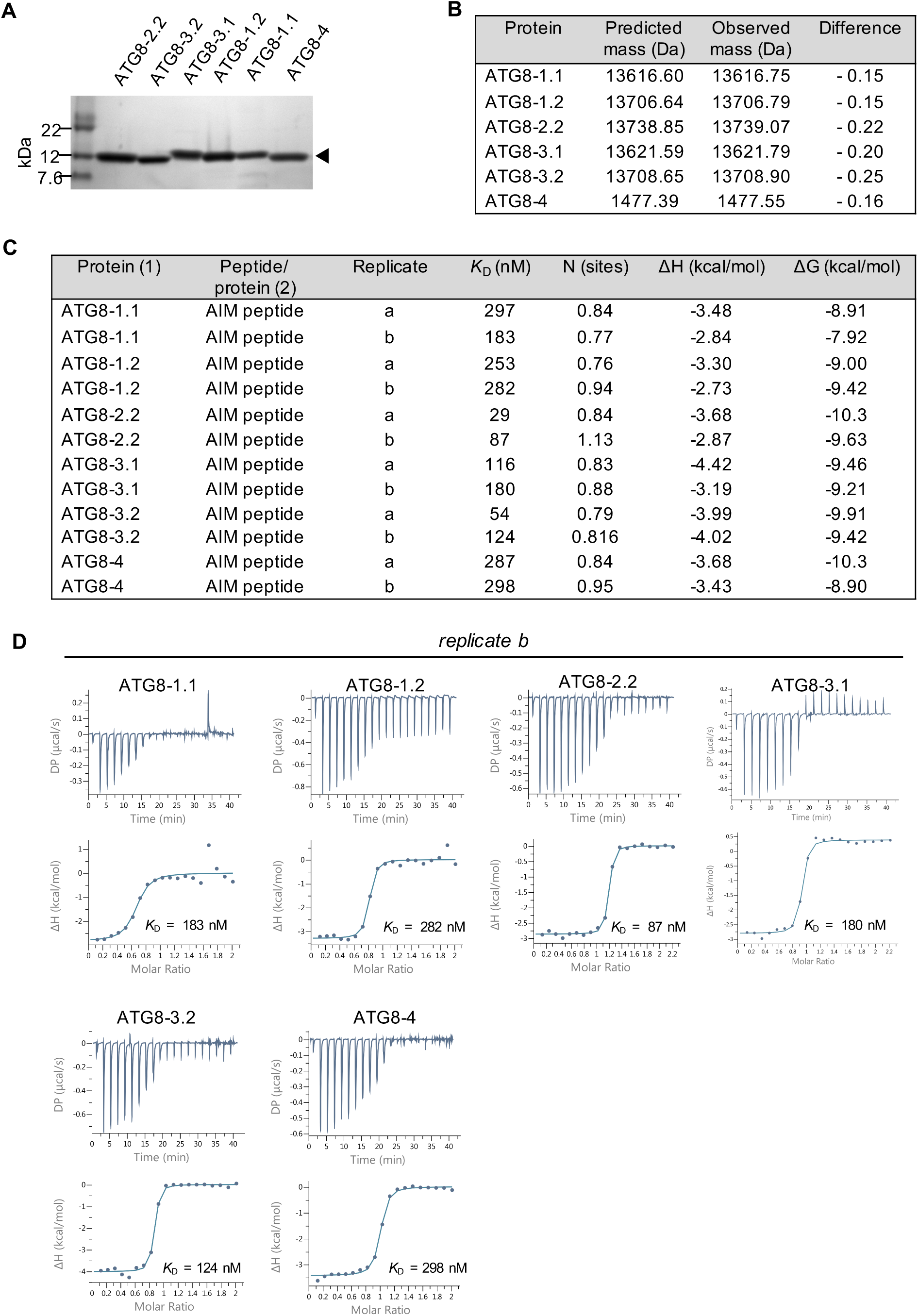
(a) Coomassie-Blue-stained SDS/PAGE gel showing purified ATG8 isoforms used in *in vitro* binding studies. (b) Intact masses for ATG8 isoforms expressed and purified in this study. (c) Table summarizing the thermodynamic and kinetic data that were extracted for each isothermal titration calorimetry (ITC) run between the PexRD54 AIM peptide and ATG8 isoforms. (d) Second replicate of ITC measuring the interaction between the PexRD54 AIM peptide and ATG8 isoforms. The top panels show heat differences upon injection of ligands and lower panels show integrated heats of injection (•) and the best fit (solid line) to a single site binding model using MicroCal PEAQ-ITC analysis software.

**Fig. S10.**
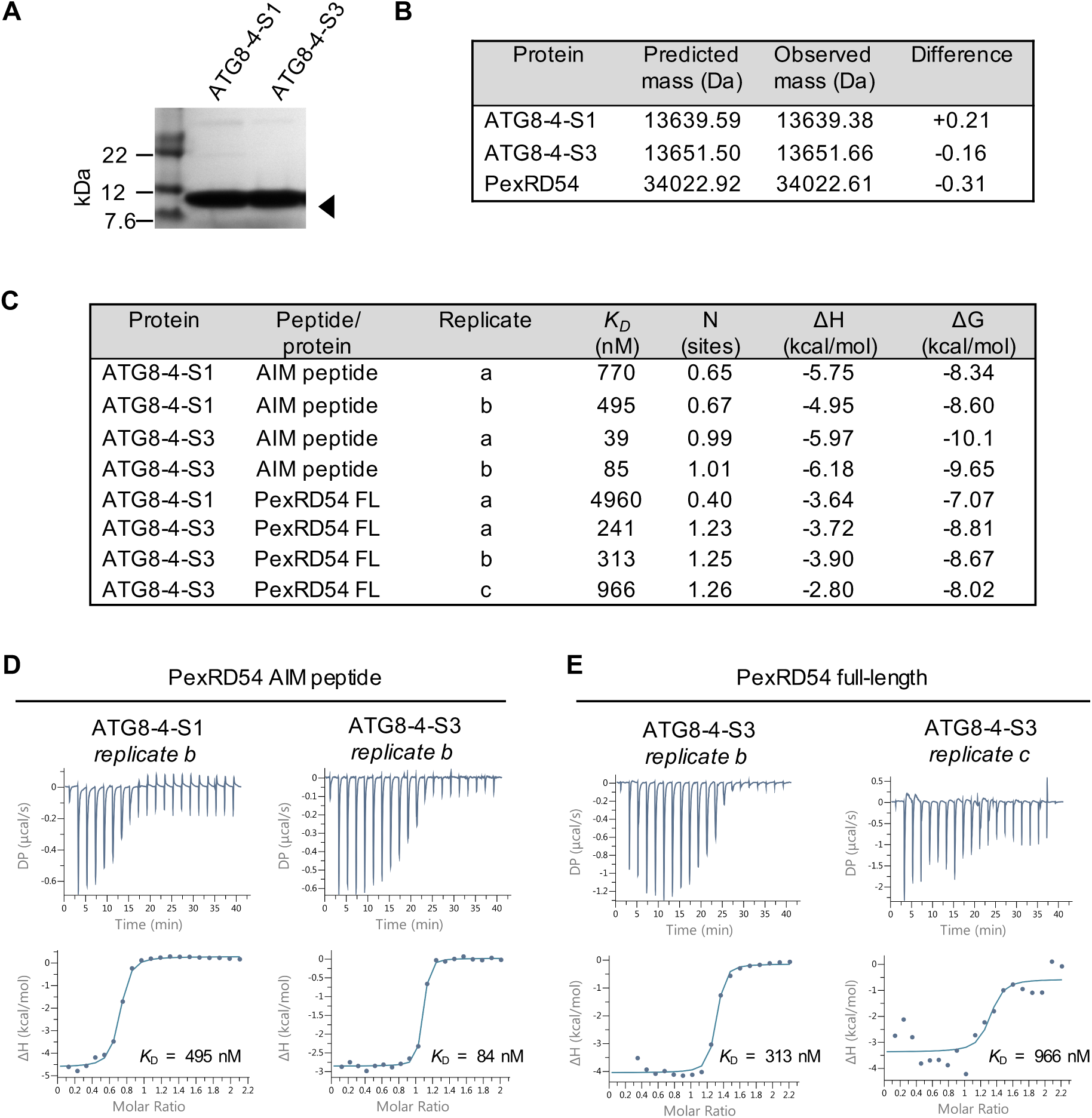
(a) Coomassie-Blue-stained SDS/PAGE gel showing purified ATG8-4-S1 and ATG8-4-S3 used in *in vitro* binding studies, (b) Intact masses for ATG8 swaps (ATG8-4-S1 and ATG8-4-S3) and PexRD54 expressed and purified in this study, (c) Table summarizing the thermodynamic and kinetic data that were extracted for each isothermal titration calorimetry (ITC) run between the PexRD54 full-length, PexRD54 AIM peptide, and ATG8 swaps, (c) Replicates of ITC measuring the interaction between ATG8 swaps and the PexRD54 AIM peptide (left) and full-length protein (right). The top panels show heat differences upon injection of ligands and lower panels show integrated heats of injection (•) and the best fit (solid line) to a single site binding model using MicroCal PEAQ-ITC analysis software.

**Fig. S11.**
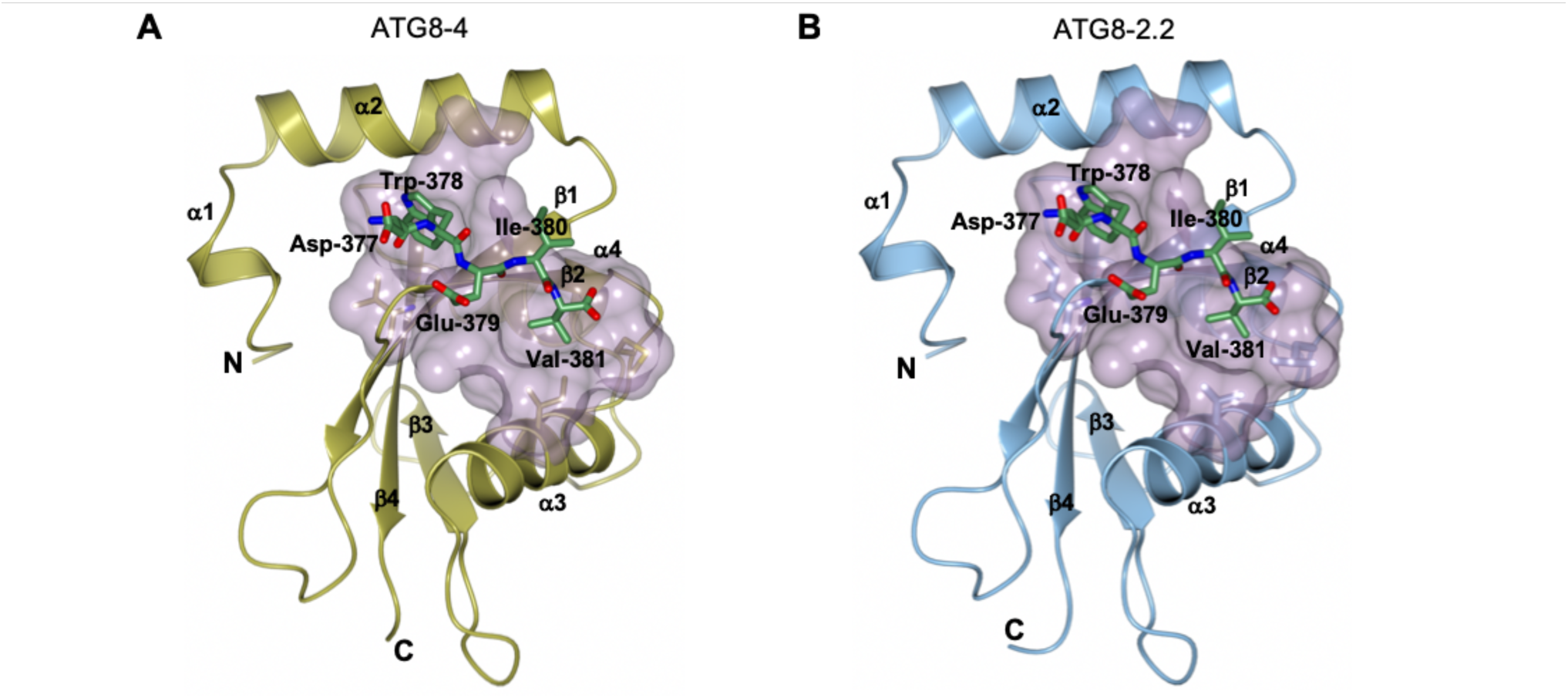
Schematic representation of (a) ATG8-4 and (b) ATG8-2.2 with PexRD54 AIM peptide in the binding cavity. The molecular surface of each ATG8 that contacts the AIM peptide is shown in magenta. The AIM peptide is shown as stick representation in each structure with residues labelled. a-helices, b-strands, N and C termini of ATG8-4 and ATG8-2.2 are labelled.

**Fig. S12.**
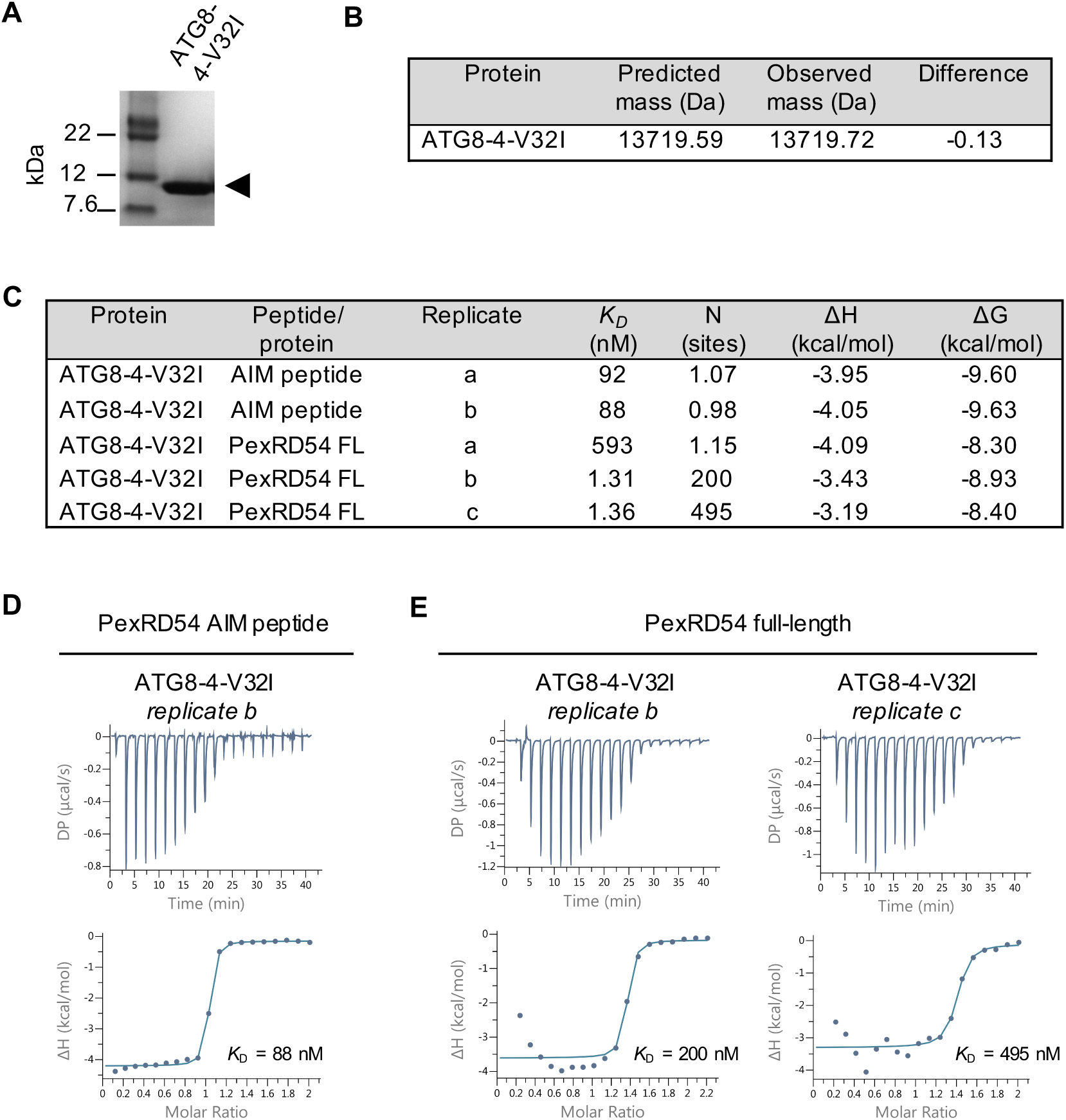
(a) Coomassie stained SDS-PAGE showing purified ATG8-4-V32I. (b) Identity of ATG8-4-V32I was confirmed by measuring intact mass using mass spectrometry. (c) Table summarizing the thermodynamic and kinetic data that were extracted for each isothermal titration calorimetry (ITC) run between the PexRD54 full-length, PexRD54 AIM peptide, and ATG8-4-V32I. (d) Second replicate of the ITC trace showing interaction between ATG8-4-V32I and PexRD54 AIM peptide. (e) Replicates of the ITC traces showing interaction between ATG8-4-V32I and the full-length PexRD54.

**Fig. S13.**
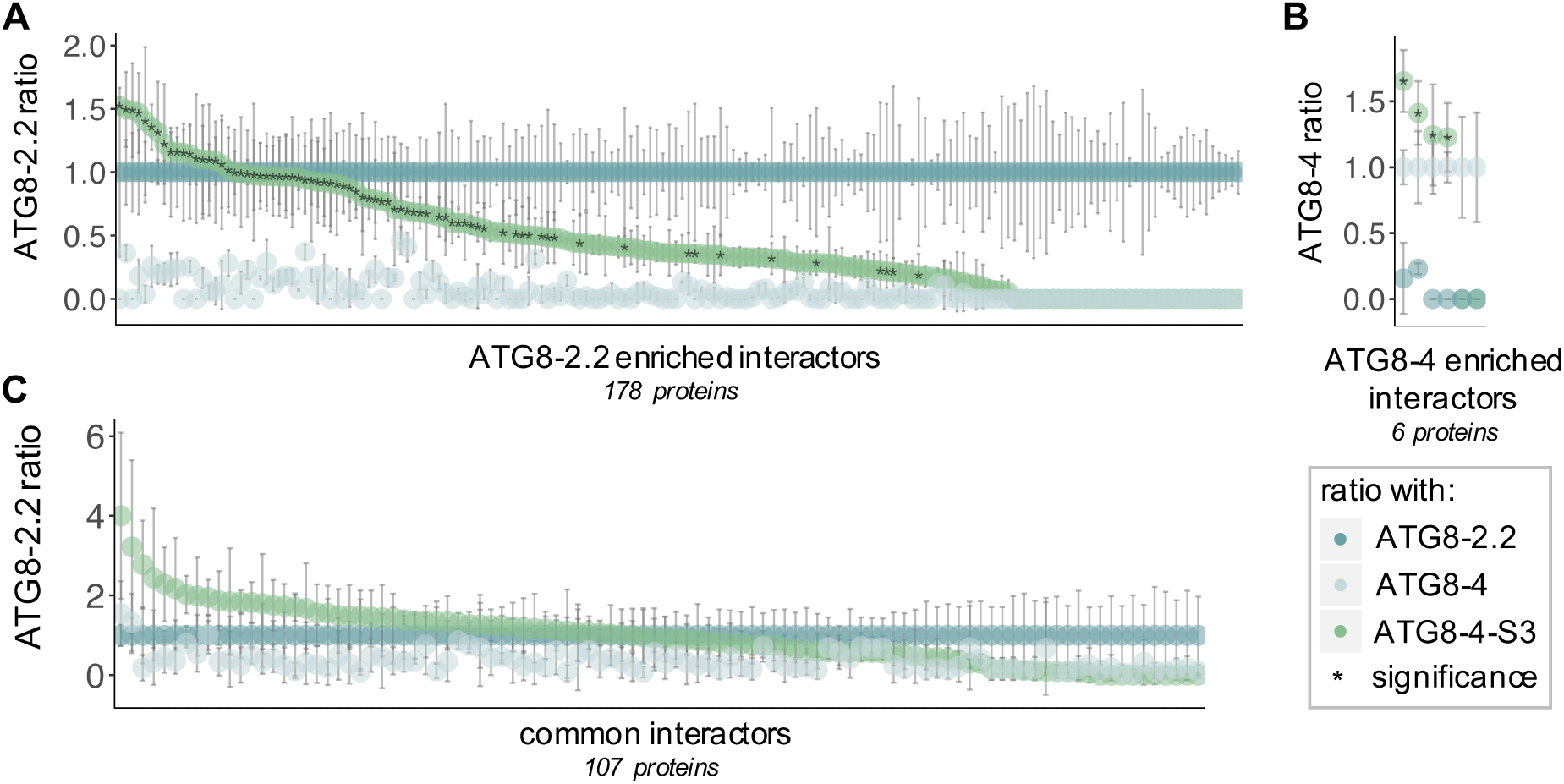
The first β-strand of ATG8 underpins interaction with plant proteins. For each interactor in the dataset, the average peptide count data for ATG8-2.2 (teal), ATG8-4 (light grey), and ATG8-4-S3 (green) were normalized to either ATG8-2.2 or ATG8-4 data based on the enrichment category being analyzed: (a) values for ATG8-2.2 enriched interactors were normalized to ATG8-2.2, (b) values for ATG8-4 enriched interactors were normalized to ATG8-4, and (c) values for common interactors were normalized to ATG8-2.2. For (a) ATG8-2.2 enriched interactors and (b) ATG8-4 enriched interactors, this highlights the difference in how ATG8-2.2 and ATG8-4 interact with each protein in the set and how the ATG8-4-S3 interactions compare. For (a) ATG8-2.2 enriched interactors, the asterisk (*) marks proteins that showed no statistical difference in their interaction with ATG8-4-S3 as compared to ATG8-2.2 (in Fig. 6d, ‘(+) S3 enrichment); for (b) ATG8-4 enriched interactors, the asterisk (*) marks proteins that showed no statistical difference in their interaction with ATG8-4-S3 as compared to ATG8-4. For (c) common interactors, the graph highlights the similarity in how ATG8-2.2, ATG8-4, and ATG8-4-S3 interact with each protein in the set; due to the lack of statistical difference, this feature is not marked.

**Fig. S14.**
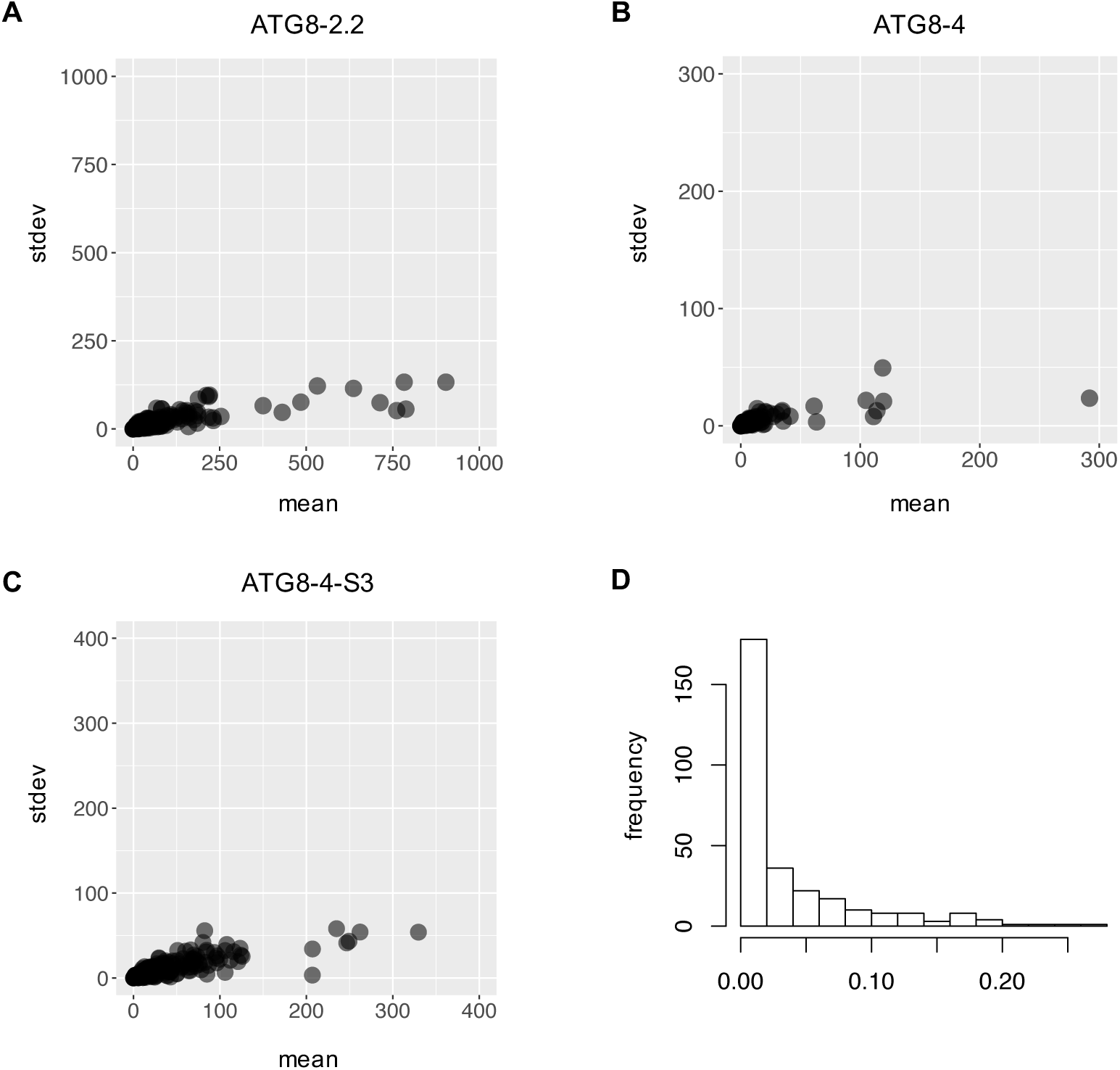
Normal distribution of comparative ATG8-4-S3 mutant analysis data. The standard deviation (stdev) versus mean is plotted for the GFP normalized peptide count data for three replicates of each construct tested in IP-MS, (a) ATG8-2.2, (b) ATG8-4, and (c) ATG8-4-S3, showing a normal distribution in each. (d) A histogram of ANOVA p-values showing the high level of significance within the dataset.

## Supplementary Tables

### File: TableS1_interactome

**Table S1**. **ATG8 interactome**. For each *N. benthamiana* interactor in the dataset (621 proteins), putative AIMs were predicted using iLIR [38], the closest *A. thaliana* and *M. polymorpha* homologs were predicted using BLAST, and the AIMs in these homologs were again predicted using iLIR. Each interactor is thus described, by column: *N. benthamiana* accession (‘Nb’), protein identification (‘Nb_protein_ID’), and number of putative AIMs (‘Nb_AIMs’); the *A. thaliana* homolog accession number (‘At’), BLAST %identity (‘At_%ID’), BLAST Expect (E) value (‘At_evalue’), protein identification (‘At_protein_ID’), number of putative *N. benthamiana* AIMs conserved (‘At_conserved_AIMs’), and gene ontology annotations (‘At_compartment’ and ‘At_process_function’) determined using Blast2GO [52]; the *M. polymorpha* homolog accession number (‘Mp’), BLAST %identity (‘Mp_%ID’), BLAST E value (‘Mp_evalue’), protein identification (‘Mp_protein_ID’), and number of putative *N. benthamiana* AIMs conserved (‘Mp_conserved_AIMs’). In addition, the average PSM values for the two replicates for each ATG8 isoform, as well as empty vector (EV) are appended.

### File: TableS2_interactome_AIM_sequences

**Table S2**. **ATG8 interactome AIM sequences and conservation.** For each *N. benthamiana* interactor in the IP-MS dataset, the putative AIMs were predicated using iLIR [38]. These putative AIMs are recorded by *N. benthamiana* accession (‘Nb’), with the start (‘Nb_start’) and end (‘Nb_end’) points of the AIM included, as well as the sequence (‘Nb_sequence’) and iLIR prediction score (‘Nb_PSSM’). For *N. benthamiana* proteins with multiple predicted AIMs, all are included. For each protein, the closest *A. thaliana* (‘At’) and *M. polymorpha* (‘Mp’) homologs were also analysed for putative AIMs, and those conserved with *N. benthamiana* were recorded. For each conserved AIM, the conservation score (number of positions conserved) was defined (‘At_conserved’ and ‘Mp_conserved’), as well as the AIM start and end sites, sequence, and prediction score.

### File: TableS3_interactome_Swap3_cross_ref

**Table S3**. **Overlap between ATG8 interactome and Swap3 interactome datasets.** The ATG8 interactome (Table S1) and Swap3 interactome (Table S4) were cross-referenced by *N. benthamiana* accession (‘Nb’) and protein description (‘Nb_protein_ID’). Interactors defined as shared between the datasets were determined by matching accession numbers (green) or in a family-based manner by exact protein description (blue).

### File: TableS4_Swap3_interactome

**Table S4. Comparative ATG8-4-S3 mutant analysis dataset.** The *N. benthamiana* proteins in the dataset (291 proteins) were divided into enrichment categories based on whether they showed a significantly (p < 0.05) stronger interaction with ATG8-2.2 or ATG8-4 as determined by an ANOVA with a post-hoc Tukey’s test; interactors that showed no significant difference in their interaction with either protein were categorized as ‘common’ (‘enrichment’ column). This resulted in 178 interactors being defined as ATG8-2.2 enriched, 6 as ATG8-4 enriched, and 107 as common. For each ATG8-2.2 enriched interactor, we determined whether ATG8-4-S3 showed a significant (p < 0.05) difference in its interaction strength compared to ATG8-2.2 using an ANOVA with a post-hoc Tukey’s test (‘S3-2.2’ column). Proteins that showed no statistical difference in their interaction with ATG8-4-S3 compared to ATG8-2.2 are categorized as ‘+’; those that showed a statistically weaker interaction are categorized as ‘-’. For each ATG8-4 specific interactor, it was determined whether ATG8-4-S3 showed a significant (p < 0.05) difference in its interaction strength compared to ATG8-4 using an ANOVA with a post-hoc Tukey’s test (‘S3-4’ column). Proteins that showed no statistical difference in their interaction with ATG8-4-S3 compared to ATG8-4 are categorized as ‘+’; those that showed a statistically weaker interaction are categorized as ‘-’. In addition to this analysis, the same interactor descriptions from the ATG8 interactome (Table S1) were included – including predicted AIMs, *A. thaliana* orthologs, GO annotations, *M. polymorpha* orthologs, and AIM conservation in both species – along with the averaged GFP normalized peptide count data for all constructs.

### File: TableS5_interactome_AIM_sequences

**Table S2**. **Swap interactome AIM sequences and conservation.** For each *N. benthamiana* interactor in the comparative ATG8-4-S3 mutant analysis, the putative AIMs were predicated using iLIR (38). These putative AIMs are recorded by *N. benthamiana* accession (‘Nb’), with the start (‘Nb_start’) and end (‘Nb_end’) points of the AIM included, as well as the sequence (‘Nb_sequence’) and iLIR prediction score (‘Nb_PSSM’). For *N. benthamiana* proteins with multiple predicted AIMs, all are included. For each protein, the closest *A. thaliana* (‘At’) and *M. polymorpha* (‘Mp’) homologs were also analysed for putative AIMs, and those conserved with *N. benthamiana* were recorded. For each conserved AIM, the conservation score (number of positions conserved) was defined (‘At_conserved’ and ‘Mp_conserved’), as well as the AIM start and end sites, sequence, and prediction score.

### File: TableS6_primers

**Table S5. Primers used in this study (5’- 3’).**

